# An experimental test of information use by North American wood ducks (*Aix sponsa*): external habitat cues, not social visual cues, influence initial nest-site selection

**DOI:** 10.1101/2020.05.08.084012

**Authors:** Elena C. Berg, John M. Eadie

## Abstract

Birds may use a variety of cues to select a nest site, including external information on habitat structure and nest site characteristics, or they may rely instead on social information obtained directly or indirectly from the actions of conspecifics. We used an experimental manipulation to determine the extent to which a California population of the wood duck (*Aix sponsa*) used social information gleaned from visual cues inside nest boxes that might indicate the quality or occupancy of that site. Over two nesting seasons, we manipulated the contents of newly installed boxes to simulate one of three states: (1) presence of wood duck eggs, indicating current use of a nest site; (2) presence of down and shell membranes, indicating a previously successful nest; and (3) control nests with fresh shavings indicating an unused box. In addition, we measured habitat characteristics of the area surrounding each box to assess the use of external, non-social information about each nest site. We found no evidence that females laid eggs preferentially, or that conspecific brood parasitism was more likely to occur, in any of the treatments. In contrast, nest site use and reproductive traits of wood ducks did vary with vegetation cover, and orientation and distance of the box from water. Our results suggest that personal information, not social information, influence initial nest site selection decisions when females are unfamiliar with a site. Social cues likely become increasingly important once nest sites develop their own history, and a population becomes well established.

**Significance Statement:** In selecting a nest site, birds may use many types of information, including habitat characteristics, their own previous breeding experience, or social cues inadvertently provided by other individuals of the same or different species. We examined information use in a Californian population of wood ducks by experimentally manipulating the visual cues within nest boxes and found that females did not use internal box cues to direct their nesting behaviors, appearing to rely on key habitat characteristics instead. These results contrast with previous studies of this system, suggesting that females may change the cues they use depending on their prior experience with a particular area. In the nest-site selection literature, there appears to be a divergence between research on passerines versus waterfowl, and we advocate unifying these perspectives.

## Introduction

Animals use various types of information to make decisions about what to eat, where to live, and how to find mates or avoid predators. They can rely on “personal” information gleaned from the physical environment or from their own private experience (i.e., trial and error), or they can use “social” information, taking advantage of signals or cues provided by other individuals of the same or different species (Danchin et al. 2004). Social information can be based on 1) inadvertent cues about another individual’s performance (typically referred to as “public” information), 2) the location of other individuals or 3) intentional signals produced by con- or heterospecifics (Danchin et al. 2004). This question of when and how animals use different kinds of information has inspired a growing number of theoretical studies and reviews (Bonnie and Earley 2007; Dall et al. 2005; Danchin et al. 2004; Dubois et al. 2012; Evans et al. 2016; Gil et al. 2019; Lee et al. 2016; Rieucau and Giraldeau 2011; Schmidt and Whelan 2010; Seppänen et al. 2007; Valone 2007), complemented by empirical studies exploring the impact of social information use on foraging behavior (Coolen et al. 2003; Machovsky-Capuska et al. 2014; Templeton and Giraldeau 1996), antipredator behavior (Frechette et al. 2014; Griffin 2004), mate choice (Nordell and Valone 1998), and breeding habitat selection (Danchin et al. 1998; Pöysä 2006; Vaclav et al. 2011). Evidence of social information use has been found in a wide array of taxa, including mammals (e.g., Ellard and Byers 2005; Lewanzik et al. 2019; Toelch et al. 2014), fish (e.g., Coolen et al. 2003; Elvidge et al. 2016; Webster and Laland 2017), amphibians and reptiles (e.g., Hobel and Christie 2016; Kar et al. 2017), insects (e.g., Avarguès-Weber et al. 2018; Grüter and Leadbeater 2014), and birds (e.g., Aparicio et al. 2007; Roy et al. 2009; Tolvanen et al. 2018).

The use of personal and social information has been especially well studied in birds, particularly in the context of nest-site selection (Campobello and Sealy 2011; Chalfoun and Schmidt 2012; Nocera and Betts 2010 and other contributors to a special issue of *The Condor*; Szymkowiak 2013). In selecting a nest site, females may use environmental cues such as food availability or habitat structure (Brown and Brown 1996; Orians and Wittenberger 1991) or nest site visibility (Bellrose and Holm 1994; Semel and Sherman 1986; Semel and Sherman 1995; Semel et al. 1988). Environmental cues are often more static or stable than social cues. Features of the habitat (e.g., physical structure, vegetation, microclimate) are unlikely to change markedly among breeding attempts and so may provide information that is reliable over multiple years. However, because success likely depends on additional dynamics operating at a local or population level (e.g., competition for nest sites, parasite loads, predation risks), static environmental cues may not always provide reliable predictors of success.

Alternatively, females may rely on information gleaned from other individuals. For example, information on nest site preferences, use, and reproductive success could be obtained by following or observing the actions of other conspecifics (Danchin et al. 1998; Pöysä 2006), heterospecifics (Mönkkönen and Forsman 2002; Parejo and Avilés 2007; Seppänen and Forsman 2007; Tolvanen et al. 2018), or both (Samplonius et al. 2017). These inadvertent socially-generated cues are frequently more ephemeral and may operate on shorter timescales, within one or just a few breeding seasons. Social cues provide immediate information on nest site use or success by other birds, but that information may not be reliable for future breeding attempts. Hence, while external habitat cues and social cues both provide useful information – the time scale and reliability of each source of information may vary. Moreover, the utility of either source of information will also depend on the history of a given resource. A newly established nest site, for example, would have little to no history and so nest site selection may be based more on external habitat cues. Conversely, as nest sites develop their own history of use and success, social cues might become more informative, albeit requiring on-going re-assessment and refinement by the user. Accordingly, animals may use different types of information at different points in the nest site selection process and as information on the quality of a site accumulates over time.

One of the most commonly explored conspecific social cues is evidence of current or past nest success (Boulinier et al. 2008; Danchin et al. 1998; Doligez et al. 2002; Kearns and Rodewald 2013; Kelly and Schmidt 2017; Parejo et al. 2008; Sergio and Penteriani 2005; Ward 2005). For example, black-legged kittiwakes (*Rissa tridactyla*) use the reproductive success of their neighbors to decide whether to emigrate (Danchin et al. 1998); a more recent study of the same species showed that individuals whose clutches failed were more likely to return to the same breeding habitat the next year if their neighbors were successful (Boulinier et al. 2008). In collared flycatchers *(Ficedula albicollis)*, immigration and emigration rates declined when reproductive success was experimentally lowered (Doligez et al. 2002). Kearns and Rodewald (2013) found that Northern cardinals (*Cardinalis cardinalis*), but not Acadian flycatchers (*Empidonax virescens*), adjusted the height and concealment of their nests in response to both personal and social information about nest predation.

Curiously, two rather distinct research trajectories have developed among researchers working on different groups of birds in their approach to investigating nest site selection. In passerines, there has been a strong behavioral ecological orientation, incorporating ideas on social information use and reliability into habitat selection models (Ahlering et al. 2010; Ahlering and Faaborg 2006; Andrews et al. 2015; Nocera and Betts 2010 and references above). Research on waterfowl and other gamebirds, in contrast, has instead focused more on evaluating external habitat cues of resource selection, specifically on the physical, environmental and resource variables birds may be tracking to hone in on appropriate nesting locations (e.g., Clark and Shutler 1999; Crabtree et al. 1989; Dyson et al. 2019; Gloutney and Clark 1997; Hines and Mitchell 1983 and see review by Eichholz and Elmberg 2014). One explanation for this is that waterfowl studies have traditionally had a more applied management focus, with an emphasis on identifying and protecting habitats that are particularly suitable for waterfowl foraging and breeding. Classic wildlife studies, like those done on many waterfowl species, have typically measured an array of relevant environmental variables without necessarily incorporating information on social behavior (but see O’Neil et al. 2014; Pöysä et al. 1998). This is not to ignore the extensive literature on physical habitat selection by passerines and other non-gamebirds (Jones 2001), but it is striking that there has been a relative paucity of research on the use of social information in the wildlife and waterfowl literature (Eichholz and Elmberg 2014; O’Neil et al. 2014; Pöysä et al. 1998).

There is an exception to this trend, specifically for species that exhibit conspecific brood parasitism (CBP), the laying of eggs in the nests of other females of the same species. CBP occurs in a wide range of taxa, including insects, fish, and birds (Andersson 1984; Brockmann 1993; Soler 2017; Yom-Tov 1980; Zink 2000), but is particularly common among waterfowl (Eadie et al. 1988; Lyon and Eadie 2008; MacWhirter 1989; Rohwer and Freeman 1989). CBP is unique in that parental and parasitic tactics coexist in the same population. Curiously, this is one area in the waterfowl literature where researchers have paid particular attention to the role of social information, possibly because CBP is inherently a social interaction among females, and the use of social cues over short time intervals may play an important role in how parasites choose among possible host nests (reviewed in Pöysä et al. 2014). Studies of social information use and CBP have been conducted on a number of waterfowl species, including common goldeneyes (*Bucephala clangula*, Dow and Fredga 1985; Pöysä 1999; Pöysä 2006), common eiders (*Somateria mollissima*, Fast et al. 2010; Lusignan et al. 2010), red-breasted mergansers (*Mergus serrator*, Thimot et al. 2020), and North American wood ducks (*Aix sponsa*, Odell and Eadie 2010; Roy et al. 2009; Semel and Sherman 1986; Semel and Sherman 1995).

In the current study we attempt to bridge the gap between these two approaches by investigating both external habitat (environmental) as well as social cues underlying nest-site selection in a California population of the North American wood duck. Previous research on this species suggests that females may rely on factors intrinsic to the site itself, preferring nest boxes in highly visible areas (Bellrose and Holm 1994; Roy-Nielsen et al. 2006; Semel and Sherman 1986; Semel and Sherman 1995; Semel et al. 1988); but see (Jansen and Bollinger 1998). Separate studies have suggested that females use social cues to assess the quality of individual nest-sites, laying preferentially in previously successful nests (Hepp and Kennamer 1992); in nests that were previously used but not necessarily successful (Roy et al. 2009); in active nests containing eggs (Clawson et al. 1979; Wilson 1993); in active nests with low numbers of eggs (Odell and Eadie 2010); or in nest boxes around which other ducks have gathered (Heusmann et al. 1980; Semel and Sherman 1986; Semel and Sherman 1995; Wilson 1993). In wood ducks, the main cause of nest failure is nest desertion, not predation, which might explain why females seem to be honing in less on previous nest success compared to the highly depredated nests of common goldeneyes (Roy et al. 2009).

To tease apart which – if any - environmental and social cues females (nesting or parasitic) may be using, we conducted an experimental field study in which we manipulated the internal social cues in newly-erected nest boxes, while concurrently collecting extensive habitat data at each nest site. We erected brand new boxes to control for previous nest use and other historical factors that might influencing nesting behavior (Pöysä et al. 2014). Wood ducks are particularly well-suited to this kind of experimental study because they readily use nest boxes and exhibit generally high levels of parasitic behavior. Over the course of two field seasons, we experimentally manipulated nest contents to mimic one of three different conditions: an unused nest (control, with wood shavings); an active nest during the laying stage (with eggs sitting on top of the shavings); or a previously successful nest (with eggshells and down). This allowed us to test whether females are using evidence of previous/current box use to direct their laying strategies, and whether these tactics differ among nesting versus parasitic females. At each nest box site, we also collected data on an array of environmental variables, including box visibility and orientation, proximity to water, and distance between boxes.

If females are preferentially selecting previously successful nest sites (i.e., “safe” sites with lower predation risk, Pöysä 1999; Pöysä 2006), then we would expect them to favor the nests with down and eggshells in them. If they are using current box use as a guide (Clawson et al. 1979; Odell and Eadie 2010; Wilson 1993), then boxes with eggs already in them should be favored – particularly by parasitic (non-incubating) females. Conversely, if females avoid nests with evidence of current occupancy, treatment boxes with eggs should be avoided. Alternatively, it is possible that females pay little or no attention to internal box cues, relying instead on key habitat characteristics that might provide more reliable long-term (static) information – at least for “new” nest sites such as these. A final possibility is that females are using some combination of personal and social information.

## Methods

### Study Area

Our study was conducted within the Putah Creek Reserve in Davis, California during March-July of 1998 and 1999. New nest boxes were erected along lower Putah Creek, located at the southern end of the Putah-Cache Creek watershed. This natural waterway winds through both urban and agricultural landscapes and is an important resource for both farmers and wildlife. Our study site was divided into two sections, 1) “Putah Creek” (PC), a 5.52 km (63.2 ha) section of the creek where a total of 37 boxes were erected 41-469 m apart (mean = 157 m; 0.62 boxes/ha), and 2) “Russell Ranch” (RR), a 1.79 km stretch (24.75 ha) located approximately 6 km downstream from PC where we erected 12 boxes at similar density, one box every 87-207 m (mean = 130 m, 0.93 boxes/ha). Along PC, 34 of the 37 boxes were erected just prior to the 1998 nesting season; 4 were erected in 1997 after the nesting season but were not set up for use until 1998. At RR, 7 of the boxes were erected in 1997 and 5 were erected at the end of the nesting season in 1998, but none of the boxes were set up for use until just prior to the 1999 nesting season. Nest box density at both sites was far lower than that reported in many other studies of wood ducks (e.g., Semel and Sherman 1995), and closely approximated natural cavity densities – e.g., 0.68 cavities/ha (Soulliere 1988), 4.0 cavities/ha, range 0.8–15.3 cavities/ha (Gilmer et al. 1978). Boxes were placed between 1.5 and 5 m (mean = 3 m) above the ground primarily on oak (*Quercus*), cottonwood (*Populus*), walnut (*Juglans*), and eucalpytus (*Eucalyptus*) trees located between 2 and 60 m (mean = 16 m) from the bank of the creek.

### Experimental Manipulation of Internal Box Environment

The visual cues influencing female nest-site selection were analyzed experimentally using the responses of females to various nesting conditions, simulated by different combinations of wood duck eggs, down, and eggshells. The responses of breeding females to nests with eggs in them (representing active or recently abandoned nests), nests with down and eggshells in them (representing either successfully hatched or predated nests), and empty (unused) nests was recorded to see which nest-site conditions most attract females. During each breeding season, we randomly assigned each nest box to equal numbers of the three treatments, defined as follows:

1. <u>Control</u>: Our control treatment consisted of a 10-cm layer of wood shavings. This is the standard way of prepping a nest box for use by wood duck females.
2. <u>Eggs</u>: To simulate an active or recently abandoned nest, we placed three eggs on top of approximately 10 cm of wood shavings. We used either unhatched wood duck eggs, wood duck eggs from a recently-abandoned nest, or when no fresh duck eggs were available, unfertilized chicken eggs. Chicken eggs are similar in color, size, and shape to wood duck eggs and thus closely approximated natural conditions.
3. <u>Down and eggshells</u>: Nesting wood ducks produce a layer of down with which they cover eggs during forays off the nest. When eggs hatch, pieces of shell and membranes are consistently left behind with the down. To simulate a successfully hatched nest, we placed a 3-cm layer of wood duck down interspersed with eggshell membranes and shell fragments onto wood shavings. The down and membranes were collected from old nests that had either produced ducklings or were predated after hatch.

In 1998, we worked only at the PC site and conducted one set of experiments, establishing treatment nests in newly-erected nest boxes between March 15 and April 24. In 1999, we included 12 boxes at the RR site and conducted two sets of replicate experiments at each site. From March 3 to March 17 before nesting began, we randomly assigned each nest box to equal numbers of the three treatments. From May 5 to May 24 we repeated the experiment and reset all boxes that had not been used and re-randomized treatments. We realize that females might have responded differently to boxes depending on their familiarity with them. For this reason, we conducted different analyses depending on each of these sets of treatments, as described in “Statistical Analyses” below.

### Nest Monitoring and Identity of Reproductive Tactics

Nest checks every other day as well as close monitoring of females determined which nest boxes were being selected by which females and whether or not the eggs laid in these boxes were subsequently incubated. During each nest check, the box was plugged to prevent the female, if present, from flushing from the nest. This minimized the danger of egg damage and allowed us to identify (and individually mark, if we had not already done so) the nesting female. We recorded the following nest stages: ‘playing house,’ (the wood shavings were disturbed, or there was a depression in the shavings, but no eggs were present) ‘laying,’ (one or more eggs were present but were at ambient temperature), and ‘incubation’ (if a female was present and eggs were warm, or if a female was absent but eggs were warm and covered with down). In active nests, we used a fine-tipped permanent marker to number the end of each egg.

We established the possible presence of parasitic females using a combination of techniques, including nest-trapping and banding of females and regular monitoring of all active nests. A nest was considered parasitized if any of three criteria were met: more than one egg was laid per day, the total number of eggs in a clutch exceeded 13, or eggs were laid after the onset of incubation (Eadie et al. 2010). These criteria are well-established proxies for nest parasitism (Brown 1984; Eadie et al. 2010; Gibbons 1986; Lyon and Eadie 2017; Yom-Tov 1980).

### External Environmental Characteristics

To examine the alternative hypothesis that females select and use nest sites based on external environmental characteristics instead of, or in addition to, internal box cues, we collected data on an extensive series of habitat features, incorporating 20 variables related to: (i) vegetation cover at the nest site, (ii) tree species and stand composition, (iii) proximity to other trees, nest boxes or water, (iv) orientation and height of the box, and (v) characteristics of the shoreline. These variables are summarized in Table 1 with abbreviations referred to in analyses.

**Table 1.** List of habitat variables measured at each nest box for experimental nests

| **Abbreviation** | **Description** |
| --- | --- |
| **Height** | Nest box height (m) |
| **Dist Nest Box** | Distance to nearest nest box (m) |
| **Comp Dir** | Compass orientation of entrance (°) |
| **Dir Water** | Compass orientation toward water from box entrance (°) |
| **Dist Water** | Distance to water from box entrance (m) |
| **% Cov Grd** | % Vegetation cover at ground - base of nest (0-100%) |
| **% Cov E** | % Vegetation cover at entrance level of nest (0-100%) |
| **% Cov 5m** | % Vegetation cover at 5 m height above nest (0-100%) |
| **Canopy HT** | Estimated height of top of tree canopy at box location |
| **% VIS** | Estimated % visibility of nest entrance (0-100%) |
| **Tree Species** | Tree species on which box was mounted: eucalyptus (EUCA), valley oak (QULO), Fremont cottonwood (POFE), California black walnut (JUCA), willow (SALX) |
| **Std Type** | Stand type – predominant tree species of stand where box mounted |
| **Std Size** | Stand size – ordinal ranking from 1 (small; 1-5 trees) to 4 (continuous) |
| **Std Density** | Stand density – ordinal ranking from 1 (sparse, 1 tree/20m) to 4 (dense; 1 tree/5m) |
| **Tree 0m** | Distance to nearest tree in front of box (0°) |
| **Tree 90m** | Distance to nearest tree to right of box (90°) |
| **Tree 180m** | Distance to nearest tree behind box (180°) |
| **Tree 270m** | Distance to nearest tree to left of box (270°) |
| **SLP** | Estimated slope from box to nearest water (°) |
| **% Shcov** | % vegetation cover along shore of nearest water |
| **TREAT** | Experimental treatment (control, down, eggs) |

We attempted to establish identical sets of experimental nest boxes strategically placed in two different habitat types, an “open” habitat in which the nest boxes are highly visible, and a “closed” habitat in which the boxes are well obscured by vegetation. This distinction between habitat types is useful because it can help determine whether females are basing their selection of nest site and their subsequent laying strategy solely on the quality of nest boxes they encounter, on their location, or on both. Some nests were occupied by other wildlife and some of the experimental treatments were disturbed (e.g. eggs broken in nest) so that sample sizes varied slightly among treatment and years.

### Statistical Analyses

Our analyses were dependent on which sets of boxes were included. Some boxes were used by other wildlife and so became unavailable (Table 2). We excluded those nests from all analyses. Further, because we used the same nest boxes erected in 1998 for experiments in 1999, a small number of boxes that were used in 1998 had previous history, creating heterogeneity in our sample. In our final analysis we excluded those nest boxes from the 1999 sample, using only the data from the first nests in 1998. Some nest sites were also used more than once by wood ducks in a given year. Again, to ensure that birds were responding only to our experimental treatment and not to additional information from the current nesting season, we analyzed only the first attempts for each box. Our goal was to ensure that, at the start of each experiment, every box had no prior history. Finally, our second replicate set of experiments in May 1999 yielded few additional nests and most boxes were unused (due to the time of the season). We therefore excluded the second replicate because it confounds the time of season when the treatments were initiated and simply adds large numbers of unused boxes that dominate the sample. Our very conservative approach has the limitation that it reduces sample size, but it ensures that we are evaluating the response of birds to the same sets of cues. Where appropriate, we note the patterns found when we relax our criteria; the results remained unchanged.

**Table 2.**
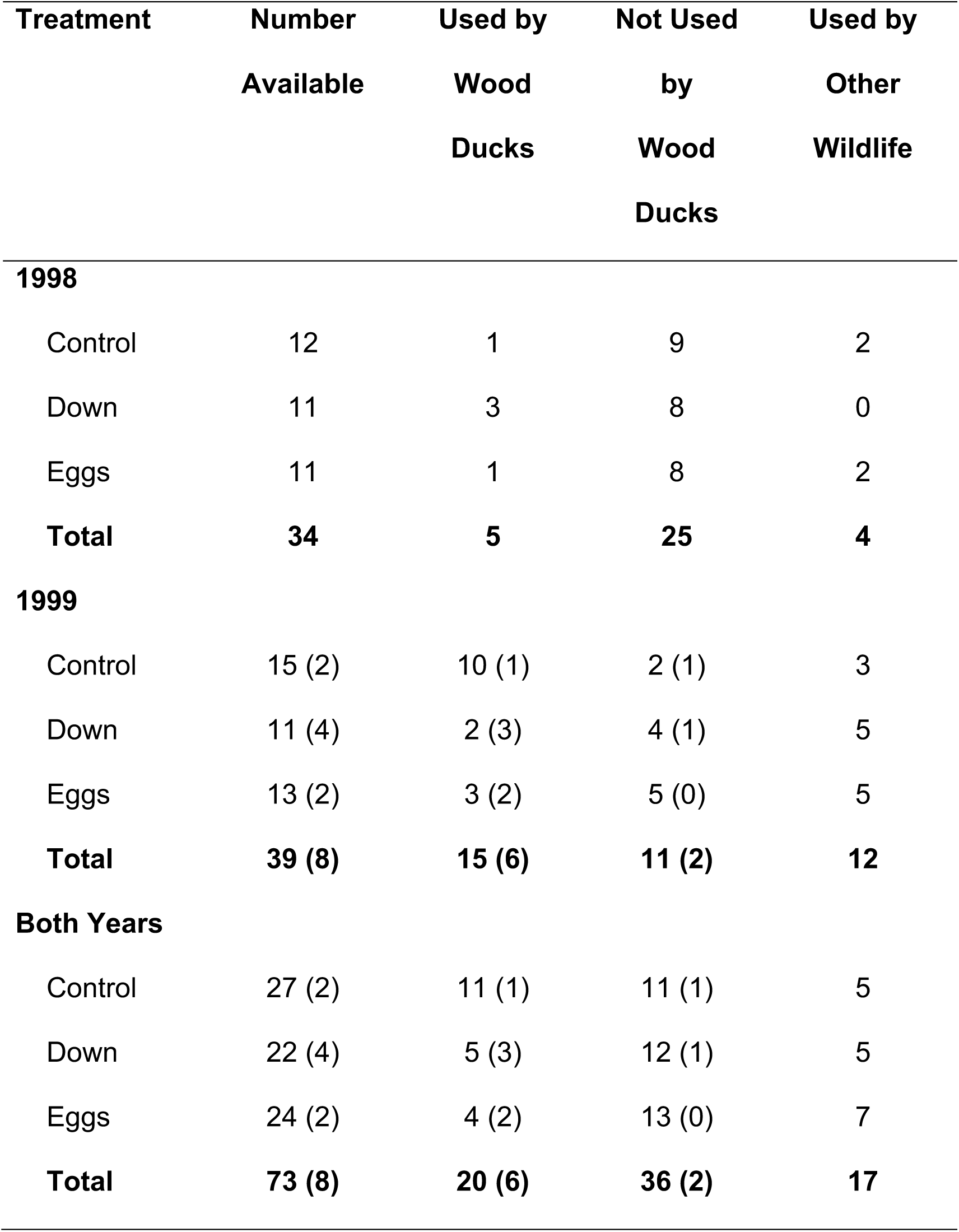
Sample sizes of experimental nests in each year. A number of nests were used by other wildlife and so were excluded from analysis. Additionally, some boxes in 1999 had been used in the previous year by wood ducks (numbers in parentheses) and so were also excluded from final analysis

| **Treatment** | **Number**<br>**Available** | **Used by**<br>**Wood**<br>**Ducks** | **Not Used**<br>**by**<br>**Wood**<br>**Ducks** | **Used by**<br>**Other**<br>**Wildlife** |
| --- | --- | --- | --- | --- |
| **1998** |  |  |  |  |
| Control | 12 | 1 | 9 | 2 |
| Down | 11 | 3 | 8 | 0 |
| Eggs | 11 | 1 | 8 | 2 |
| **Total** | **34** | **5** | **25** | **4** |
| **1999** |  |  |  |  |
| Control | 15 (2) | 10 (1) | 2 (1) | 3 |
| Down | 11 (4) | 2 (3) | 4 (1) | 5 |
| Eggs | 13 (2) | 3 (2) | 5 (0) | 5 |
| **Total** | **39 (8)** | **15 (6)** | **11 (2)** | **12** |
| **Both Years** |  |  |  |  |
| Control | 27 (2) | 11 (1) | 11 (1) | 5 |
| Down | 22 (4) | 5 (3) | 12 (1) | 5 |
| Eggs | 24 (2) | 4 (2) | 13 (0) | 7 |
| **Total** | **73 (8)** | **20 (6)** | **36 (2)** | **17** |

We used generalized linear models (GLMs) throughout (JMP 2018). Our first set of analyses focused on nest use as influenced by our experimental treatments; given strong year effects we include both treatment and year in all models. We examined several response variables; we first considered Use/No Use of the nest box as indicated by the presence of eggs and/or an incubating female. For sites that were used, we then examined the Number of Eggs Laid, Number of Eggs Hatching, and Date First Egg laid (Julian). Use/No Use was a binomial response variable, and we examined the effects of the experimental treatments using a GLM with a binomial distribution and logit link function. Number of Eggs Laid, Number of Eggs Hatched, and Date of First Egg Laid were treated as count data, and we examined the effects of the experimental treatments using a GLM with a Poisson distribution and a log link function. GLMs and Poisson models are prone to overdispersion and without correcting can give erroneous results (JMP 2018). Accordingly, for each analysis we included an overdispersion correction parameter (Pearson Chi-square deviance divided by the degrees of freedom (DF) for the full sample in the model). With overdispersion, a correction will be more robust and possibly more conservative. We employed the Firth bias-adjusted method to fit the model (Firth 1993).

Our second set of analyses examined a number of external habitat and nest characteristics that might provide an alternative source of information for breeding females. We initially considered 20 external habitat and nest variables (Table 1). A principal components analysis did not help to reduce these to a simple interpretable set of habitat dimensions, and so we retained the original variables. As in the first set of analyses, we examined several response variables, first considering Use/No Use of the nest site; for sites that were used, we then examined the effect of habitat characteristics on the Number of Eggs Laid, Number of Eggs Hatching, and Date First Egg laid (Julian). To explore the potential influence of these variables, we initially conducted simple bivariate analysis to identify a smaller subset of external characteristics that might influence each response variable, and we conducted exploratory stepwise selection methods (forward selection using minimum AICc as guiding rule). From these analyses, we focused on the subset of habitat variables that appeared to have some influence on each response variable. We conducted GLMs as described above, fitting multiple models with different combinations of habitat variables. We also contrasted each model with a similar model that included experimental treatment as an additional factor. Although many of these models were highly significant in a statistical sense (P- values << 0.01) we instead ranked models used a model selection approach based on minimum values of AIC_c_ (Akaike Information Criterion corrected for small sample size; Burnham and Anderson 2002). We emphasize that the goal of these analyses was *not* to fit a predictive model, nor even to determine which habitat variables might be best predictive of wood duck nest use. Rather, we were most interested in determining: (1) whether any subset of external variables had utility in evaluating wood duck use (do external cues matter?) and more importantly, (2) whether internal nest cue treatments provided additional or better explanatory power.

## Results

A total of 34 treatments was established in 1998, and 47 treatments in 1999 (Table 2). However, eight boxes in 1999 had been used by wood ducks in the previous year or in a previous attempt in 1999 and were thus excluded from the final analysis. The second replicate set of treatments in boxes in late May 1999 (N = 32) was also excluded to avoid confounding date and lack of use late in the season (see Methods). Sample sizes differed slightly between years and treatments due to use or interference by other wildlife (Table 2).

Manipulation of the internal box environment through the experimental addition of either eggs, egg shells and down, or wood shavings had little effect on box use by wood duck females. Over the 1998 and 1999 breeding seasons, females nested in 20 of the 81 boxes that were included in the experiment (Table 2). After excluding boxes used previously by wood ducks and those used by other wildlife, 56 Treatment boxes were available for analysis (22 Control, 17 Down, 17 Egg treatments). We found no effect of Treatment on Nest Box Use (Likelihood ratio X^2^ = 2.75, P = 0.25; Table 3). Contrary to our predictions, there was a trend for Control boxes to be used more (50%) compared to either Down (29%) or Egg treatments (24%; Figure 1). This pattern did not change when we relaxed our criteria and included boxes that had been previously used (Control 50% of 24 boxes used, Down 38% of 21 boxes used, and Egg 32% of 19 boxes used). Year had a strong influence on Nest Box Use (Likelihood ratio X^2^ = 9.95, P = 0.002; Table 3), largely because nest box use was low in 1998 compared to 1999 (Table 2).

**Table 3.**
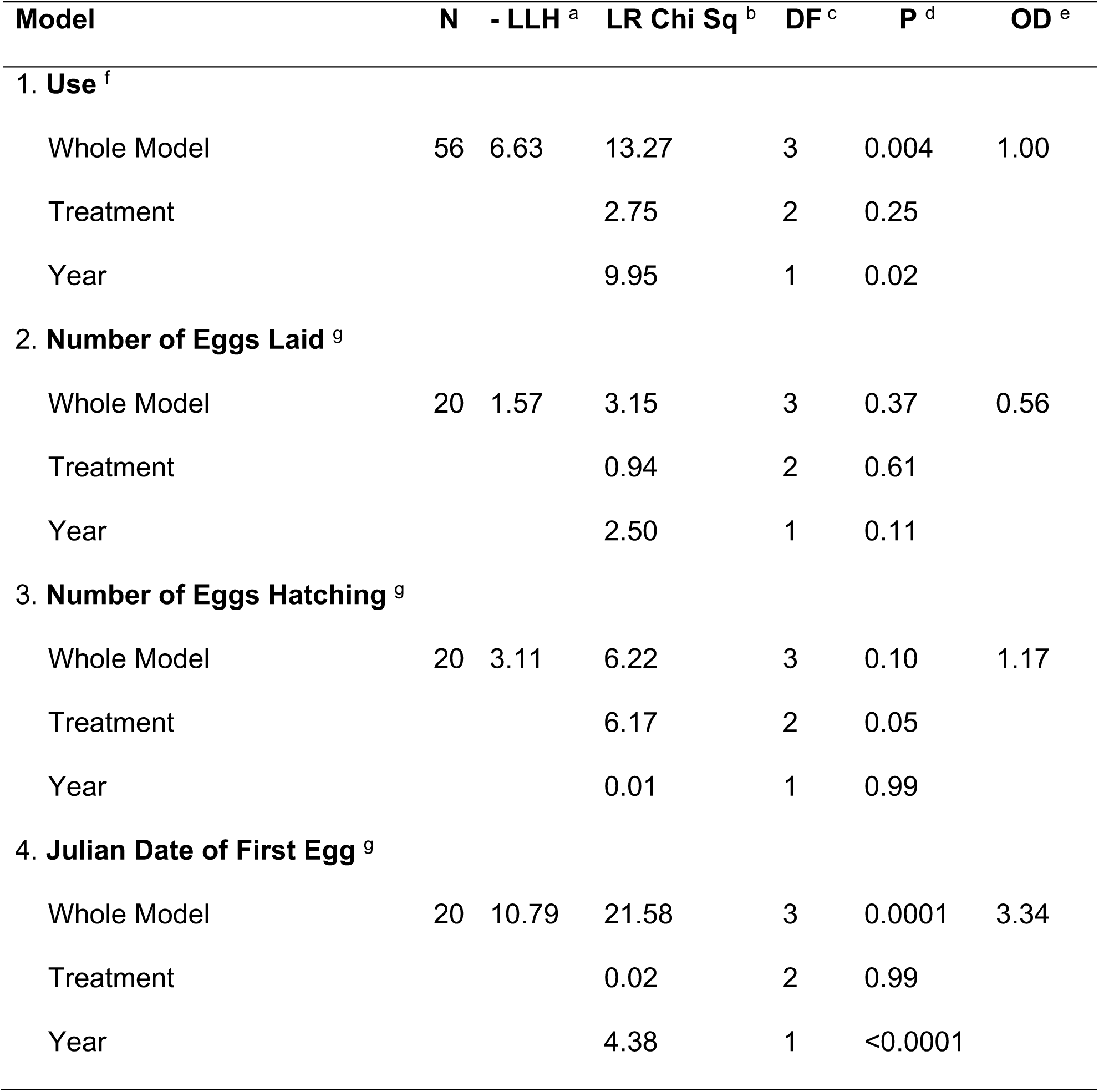
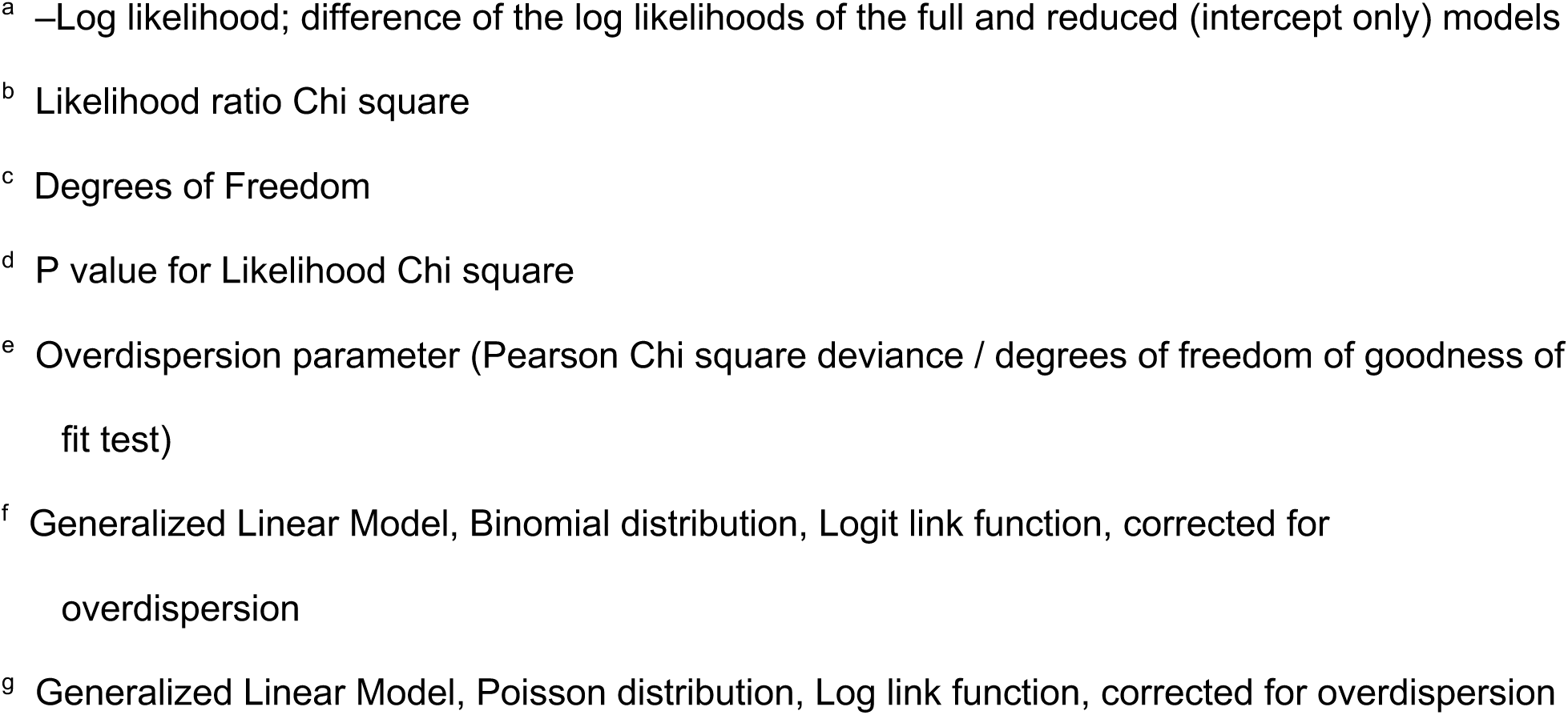
Generalized Linear Models (GLM) to examine the influence of nest box treatment (control, down added, eggs added) and year (1998, 1999) on nest box use, number of eggs laid, number of eggs hatching, and date of first egg laid for wood ducks near Davis CA. Analyses were conducted for only the first nest attempts at each box each year

| Model | N | - LLH <sup>a</sup> | LR Chi Sq <sup>b</sup> | DF <sup>c</sup> | P <sup>d</sup> | OD <sup>e</sup> |
| --- | --- | --- | --- | --- | --- | --- |
| **1. Use <sup>f</sup>** |  |  |  |  |  |  |
| Whole Model | 56 | 6.63 | 13.27 | 3 | 0.004 | 1.00 |
| Treatment |  |  | 2.75 | 2 | 0.25 |  |
| Year |  |  | 9.95 | 1 | 0.02 |  |
| **2. Number of Eggs Laid <sup>g</sup>** |  |  |  |  |  |  |
| Whole Model | 20 | 1.57 | 3.15 | 3 | 0.37 | 0.56 |
| Treatment |  |  | 0.94 | 2 | 0.61 |  |
| Year |  |  | 2.50 | 1 | 0.11 |  |
| **3. Number of Eggs Hatching <sup>g</sup>** |  |  |  |  |  |  |
| Whole Model | 20 | 3.11 | 6.22 | 3 | 0.10 | 1.17 |
| Treatment |  |  | 6.17 | 2 | 0.05 |  |
| Year |  |  | 0.01 | 1 | 0.99 |  |
| **4. Julian Date of First Egg <sup>g</sup>** |  |  |  |  |  |  |
| Whole Model | 20 | 10.79 | 21.58 | 3 | 0.0001 | 3.34 |
| Treatment |  |  | 0.02 | 2 | 0.99 |  |
| Year |  |  | 4.38 | 1 | <0.0001 |  |

<sup>a</sup> –Log likelihood; difference of the log likelihoods of the full and reduced (intercept only) models <sup>b</sup> Likelihood ratio Chi square <sup>c</sup> Degrees of Freedom <sup>d</sup> P value for Likelihood Chi square <sup>e</sup> Overdispersion parameter (Pearson Chi square deviance / degrees of freedom of goodness of fit test) <sup>f</sup> Generalized Linear Model, Binomial distribution, Logit link function, corrected for overdispersion <sup>g</sup> Generalized Linear Model, Poisson distribution, Log link function, corrected for overdispersion

**Fig 1.**
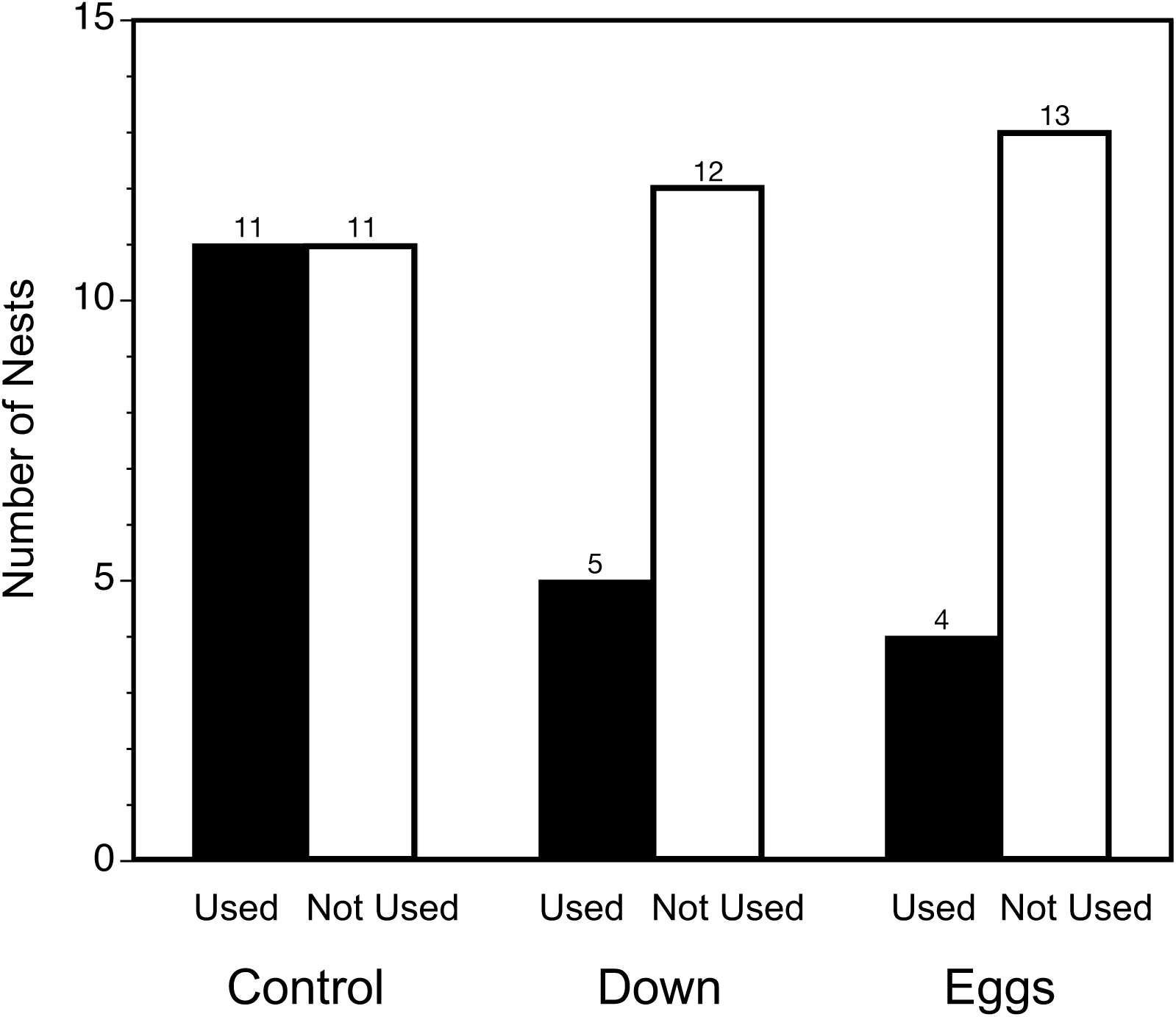
Use of new nest boxes by wood ducks in California in response to internal visual social cues. New boxes contained down and eggs shells indicating a previously successful hatch (Down), eggs without down indicating current use (Eggs), or shavings indicating no current or previous use (Control). Used boxes are shown by solid black bars, unused boxes are shown by open bars, numbers above provide the number of nests in each category.

We also found no influence of treatment on Number of Eggs Laid or Julian Date of First Egg (Table 3) although there was a significant effect of treatment on the Number of Eggs Hatching (Likelihood ratio X^2^ = 6.17, P = 0.05; Table 3). Fewer eggs hatched in the Egg treatment (Figure 2). The Julian date of First Egg was influenced by year; nesting was considerably earlier in 1999.

**Fig 2.**
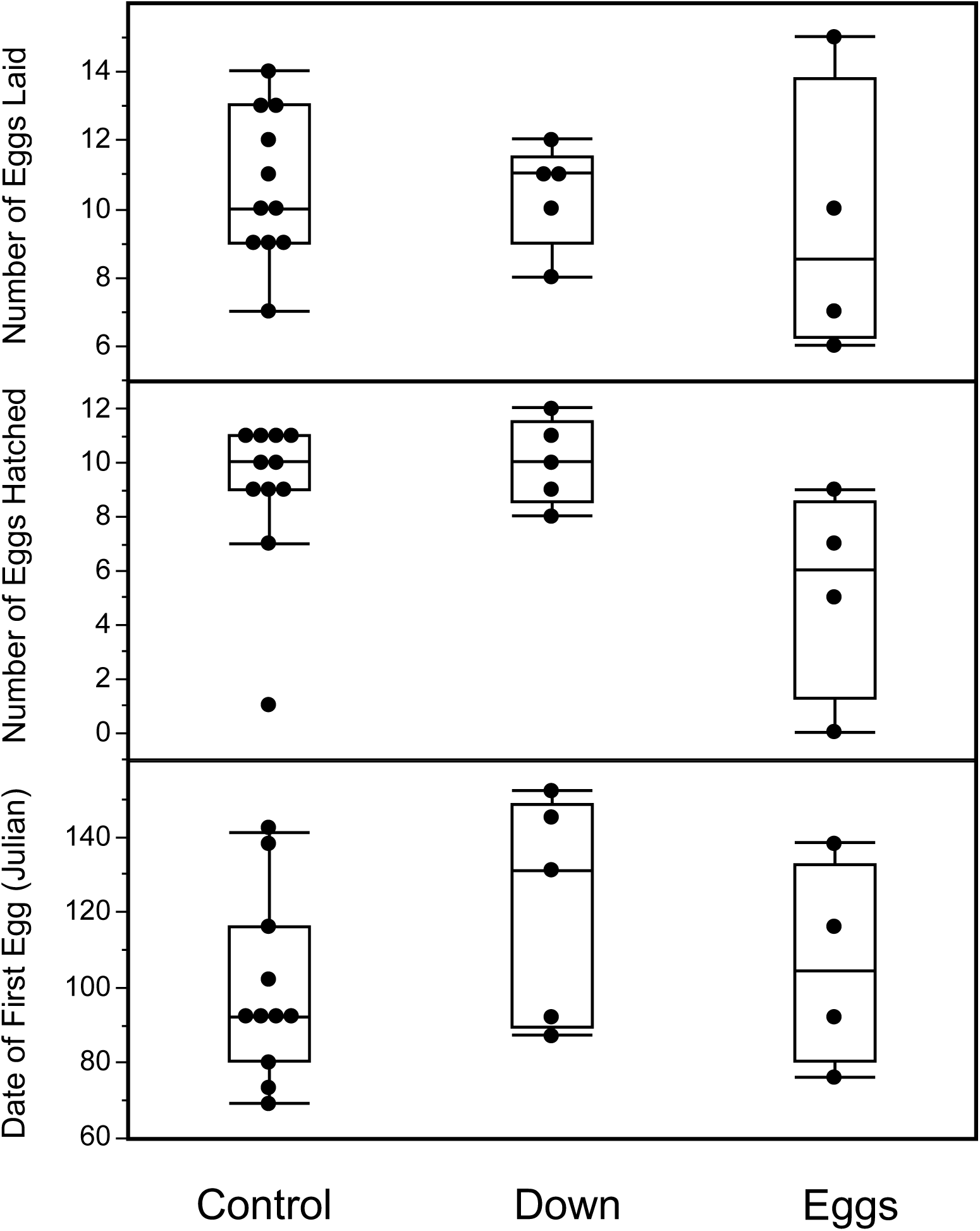
Measures of reproductive success in the three nest box treatment groups (Control, Down and Eggs). Solid points are values for individual nests with box plots showing the median, 25^th^ and 75^th^ quantiles (box), and range excluding outliers (vertical line). Top: Number of eggs laid in each nest, Middle: Number of eggs that hatched in each nest, and Bottom: Julian date of first egg laid in each nest.

Our analysis of a comprehensive set of external habitat cues suggest that a small number of characteristics influence nest use and reproductive success of wood ducks (Table 4). Notably, nest box Use was influenced by nest box visibility, presence of trees in front of the box, and orientation. Both the Number of Eggs Laid and the Number of Eggs Hatching were influenced by the direction and distance to water (Table 4). Several of these patterns were highly significant (P< 0.01), indicating that external habitat characteristics influence nest site use and investment in newly erected nest boxes.

**Table 4.**
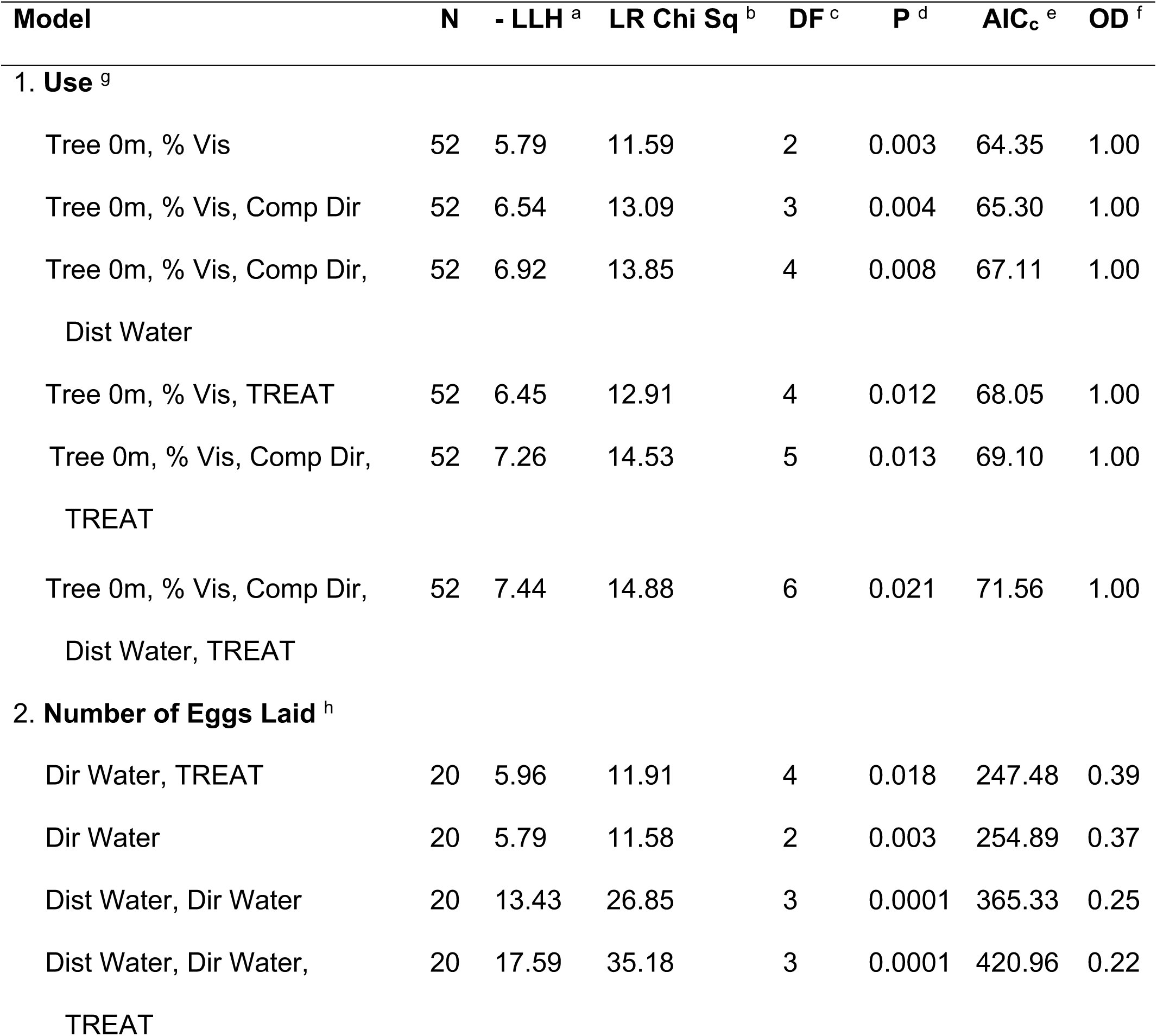
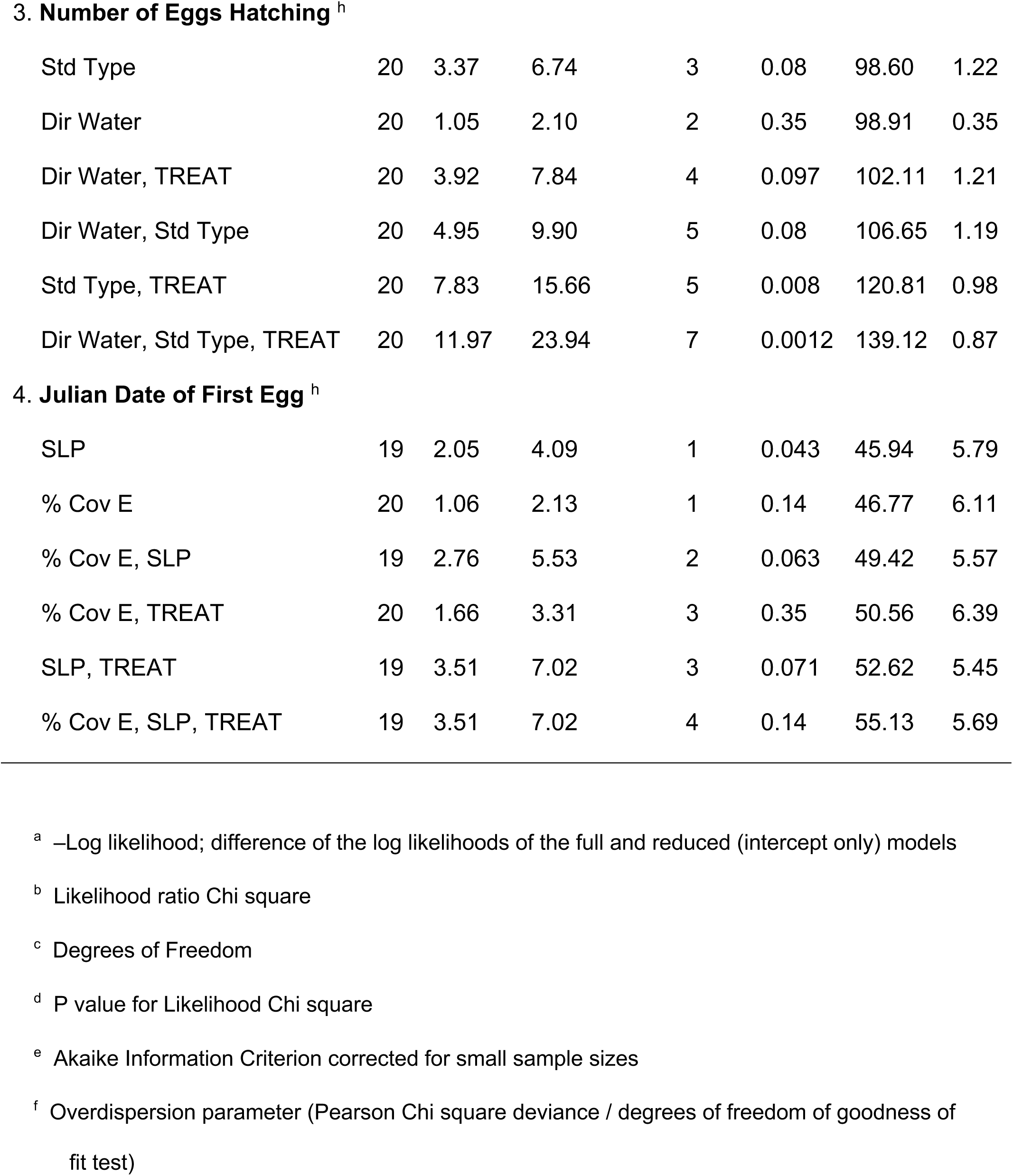
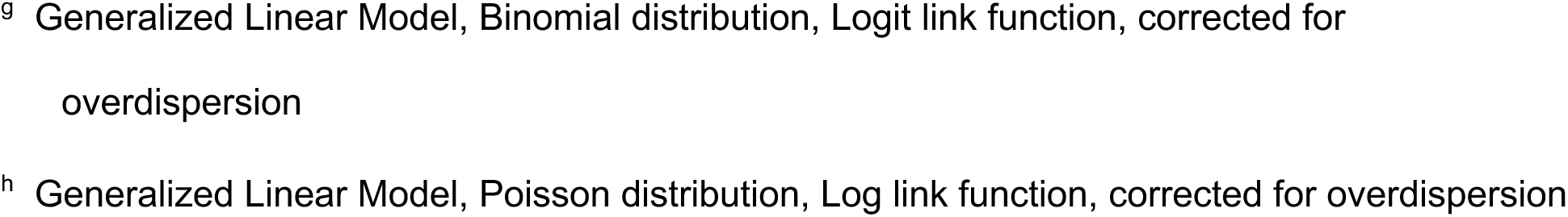
Generalized Linear Models (GLM) to examine the influence of Habitat Variables and Treatment on nest box use, number of eggs laid, number of eggs hatching, and date of first egg laid for wood ducks near Davis CA. Data for both years are included in analyses. Models are ranked using AIC_c_ Akaike Information Criterion. Habitat variable abbreviation and description are in Table 1. TREAT: experimental nest treatment (control, eggs, down)

A central question for our study was how wood ducks might make use of two different kinds of information: external habitat cues versus social cues. To fully contrast these alternatives, we considered the habitat models identified above (Table 4) and ran the models with the same subset of habitat variables but also including the effects of experimental treatment to allow both sets of ‘cues’ to compete for the data. These analyses further suggest that wood ducks do not make use of social cues but instead rely on a few habitat characteristics. For three of the four reproductive measures considered, only habitat variables were included in the top models (ΔAic_c_ > 3 or greater, Table 4). The one exception was for the Number of Eggs Laid, for which inclusion of treatment effects in addition to direction to water comprised the top model (ΔAic_c_ = 7.4, Table 4).

Finally, we found no evidence that any of the treatment nests were more likely to be parasitized. Considering all treatment nests (including those previously used) there was evidence of only 6 nests being parasitized as indicated by observations of > 1 female on the nest, clutch size sizes >13 eggs, or eggs laid after incubation. Of these, 2 parasitized nests occurred in each treatment.

## Discussion

Over the course of our two-year field experiment, we found no evidence that wood duck females used social cues about the internal state of the nest box at the beginning of the breeding season, at least not when boxes were brand new to an area. This result was somewhat surprising given that a recent study of long-term patterns of nest use by wood ducks on our study area (including Putah Creek although not during the years of the current study) found that females were more likely to use nest sites that had been previously used by other females (Roy et al. 2009). Other studies likewise suggest that wood ducks use social cues to assess the quality of individual nest-sites, such as evidence of previous success (Hepp and Kennamer 1992); nests containing eggs (Clawson et al. 1979; Odell and Eadie 2010; Wilson 1993); or nest boxes around which other ducks have gathered (Heusmann et al. 1980; Semel and Sherman 1986; Semel and Sherman 1995; Wilson 1993). In our previous study (Roy et al. 2009), we found no evidence that previous success influenced future nest box use, and so we hypothesized that females were cueing in to activities or signs of use by other females. If this were the case, and females were using internal box cues rather than actual knowledge of box use to guide their decisions, there should have been no difference in box use between egg treatments (representing active nest) and shells-and-down treatments (representing a previously successful nest), but significantly more use of those two treatments combined than of unused boxes (those with wood shavings only). In fact, although we did find that females were just as likely to use boxes with eggs and shells-and-down, if anything they showed a slight preference for empty “unused” nests with just shavings inside. Our results suggest that internal cues do not attract females to a nest-site and may even cause females to avoid these sites.

Although females were not using social cues about previous or current box use to choose a nest-site, they did seem to be paying attention to specific habitat characteristics. They were significantly more likely to nest in boxes that were located in more open visible locations, with few trees in front of the entrance and oriented towards and closer to water (Table 4). None of these patterns are unexpected and suggest that, for the females in our experiment at least, simply finding a nest site may be the central challenge and external habitat features might be most influential.

In stark contrast to our study, evidence for social information use has been documented in a number of other waterfowl species, particularly in the context of conspecific brood parasitism. For example, in cavity-nesting common goldeneyes, parasitic females appear to be using predation risk as a cue, preferentially selecting nest boxes that have not been depredated the previous year (Pöysä 1999; Pöysä 2006; Pöysä et al. 2010; Pöysä and Paasivaara 2015). In contrast, a study of red-breasted mergansers in Canada (Thimot et al. 2020) found that females were not preferentially selecting “safe” nest sites, likely because egg predation rates were low in this population. Rather, the presence of conspecifics seemed to be a cue: artificial nests with no host attracted fewer brood parasites (Thimot et al. 2020). Among common eiders, also a ground-nester, nest visibility impacts parasitism rates more than nest site safety (Lusignan et al. 2010). Another experimental study of eiders examined the specific cues females might use, indicating that they are more likely to lay in nests that had nest materials (down) in them – indicating previous nest success (Fast et al. 2010). Finally, mallards (*Anas platyrhynchos*, Pöysä et al. 1998) and lesser scaup (*Aythya affinis*, O’Neil et al. 2014) apparently use social cues such as conspecific density and proximity to previously successful nesting habitats in selection of breeding and nesting sites.

The question thus remains: why didn’t wood ducks in our study use social cues to guide their nesting choices, despite evidence from other years, populations and species suggesting that such social cues may be informative? We suggest several possibilities. *First, it may be that our experimental design and/or sample sizes were insufficient to detect patterns that might exist*. Although we erected a large number of nest boxes and established 81 possible treatment nests over two years, the final sample size was dictated by how the birds responded. As expected, there was little use in the first year but considerably more in the second (accounting for the strong year effect in nest box use; Table 3). Moreover, by using brand new boxes, our goal was to eliminate any previous history associated with each box and so, by design, we knew it would take time for birds to discover and use our nest boxes. Indeed, it was this very process of initial nest site selection that we wished to explore to determine how external habitat cues relative to internal visual social cues might influence nest site selection decisions by new females. Further, we used very conservative criteria for inclusion in our analyses, removing any site that was used in the first year (even though it was a new box that year) from analysis in the second year to ensure that previous history would not confound our analyses. Nonetheless, even when we relaxed these strict criteria, the same patterns remained – females did not disproportionately use nests with evidence of prior use. In fact, if anything, the trend was in the opposite direction regardless of which boxes were included – control boxes were more likely to be used, albeit not significantly (Figure 1), and huge sample sizes and a strong shift in patterns of nest site use would be required to alter the results. Finally, despite the more restricted samples sizes when we applied our conservative criteria, we were still able to detect statistically significant differences when considering habitat variables (Table 4). We conclude that experimental design or restricted sample sizes cannot account for the lack of use of visual social cues by wood ducks in our study; the patterns appear to be robust.

*A second possibility is that internal box cues simply are not used by these wood ducks to assess nest quality, at least during a female’s first nest attempt of the season*. If cavities are a rare commodity in nature, simply finding a nest that is usable may be the top priority. Alternatively, the lack of attention to internal cues may have more to do with when and how females prospect for nests. Common goldeneyes, which do appear to use box cues (Pöysä 2006), prospect for nests at the end of the breeding season (Eadie and Gauthier 1985; Eadie et al. 1995), when evidence of recent nesting activity is presumably still fresh. In contrast, wood ducks prospect for nests in the spring (Bellrose and Holm 1994; Dixon 1924; Hepp and Bellrose 1995). If the timing of nest searching is key, and given that wood duck females do not regularly encounter evidence of previous nesting activity in the spring, selection might not have acted on females to recognize or respond to internal box cues. Also, female wood ducks in general may be less selective than goldeneyes in part because of the relatively low rates of nest predation in wood ducks (discussed in Roy et al. 2009), and/or the speed with which wood ducks reach reproductive maturity. Female wood ducks reproduce at one year of age (Bellrose and Holm 1994), whereas goldeneyes exhibit deferred maturity and typically do not breed until they are two years of age or older, increasing the opportunity for nest exploration and information use (Eadie and Gauthier 1985; Eadie et al. 1995). However, this would not explain why other populations of wood ducks (Clawson et al. 1979; Hepp and Kennamer 1992; Wilson 1993), and even the same population of wood ducks (Odell and Eadie 2010; Roy et al. 2009) do seem to be paying attention to box cues such as the presence of eggs or down.

*A third possibility is that female wood ducks do rely on social cues, but pay more attention to the presence or activity of other wood duck females*. Betts et al. (2008) refer to such cues as ‘location cues’ indicated by the presence or position of other individuals, in contrast to ‘public information’ indicated by the success or performance of other individuals at the site (Danchin et al. 2004; Valone 1989). For example, a number of studies of other wood duck populations have suggested that females may be using cues such as the presence of females at the nest (Heusmann et al. 1980; Semel and Sherman 1986; Semel and Sherman 1995; Wilson 1993). When Wilson (1993) placed decoys of females near nest boxes, brood parasitism rates at those nests increased. This decoy effect has also been found in a number of other non-waterfowl bird species. For example, in obligate brood parasitic species such as cuckoos, simply placing experimental parasitic eggs in host nests did not elicit the maximum response by the host; the presence of cuckoo females nearby or at the nest (or a taxidermic mount) significantly increased rejection rates of experimental eggs (Davies and Brooke 1988; Langmore et al. 2009; Moksnes et al. 1993). This suggests that physical cues such as the presence of eggs alone may be insufficient to elicit a behavioral response; it may not be evidence of use, but rather visual confirmation of active use that matters. More recent data for our population of wood ducks also points to the strong influence of social information use and conspecific activity. For the past six years we have used Passive Integrated Transponders (PIT tags) and radio frequency identification detection (RFID) readers on every nest box on over 200 boxes at four study sites and we have PIT-tagged over 500 females. These data have revealed surprising and remarkable evidence that females prospect for nests in groups, visit a large number of nest sites before breeding, and that different sites – even close by – attract very different numbers of females, suggesting that conspecific cueing and information use may yet play a significant role in nest site selection processes by wood ducks (JME and colleagues, unpublished data).

*A final intriguing possibility is that females use different kinds of information, including both social and environmental cues, but their relative use of these cues varies over space and time and may be sequentially applied*. Our specific findings (use of habitat cues, not internal box cues) could be explained by the fact that our nest boxes were new. In the absence of information on past history of the boxes, females might instead utilize external habitat characteristics as the best initial estimate of the quality of the nest site. The challenge for a newly breeding female is simply to find a relatively rare but suitable nest site, and habitat cues would be available and perhaps more predictable than social cues. A shift to reliance on social cues may come only after more information about a site is acquired and the site has developed its own history of use and success. External habitat cues, over time, may not adequately predict local dynamics such as the influence of local densities of competitors (conspecifics and other species), predators, or the buildup of ectoparasite loads of lice, fleas, or mites in a nest box. At a new breeding site, such as the ones we established in 1998-1999, females may first need to gain familiarity with the “real estate” in the neighborhood, before shifting their attention to the activities and success of their neighbors. As nest sites develop a history, more refined assessments of nest site quality are possible, and it is here when social cues may be most useful. Thus, it is not so much a question of *do* birds use *either* external personal (habitat cues) *or* social information cues, as often posed, but rather *when* and under what circumstances might either or both types of cues be useful. This could also account for the observation that different studies, even on the same species such as wood ducks, yield different results regarding information use (see above). With over 20 years of data on this population, and six years of PIT tag and RFID data, we should be able to address this possible shift in focus in future studies.

We also found no evidence that parasitic and parental females differed in box use, although the frequency of parasitism was low during the two years of our study. This again may be a consequence of the early stage of our study, such that local populations had not yet increased and competition for nest sites was low; conspecific parasitism may be less frequent under these conditions (see Semel and Sherman 1986; Semel and Sherman 1995; Semel et al. 1988). There is evidence that CBP in wood ducks is density-dependent (Clawson et al. 1979; Heusmann et al. 1980; Semel and Sherman 1986) and so it would not be unexpected that parasitic females might rely more on the use of social cues to select host nests in larger established populations as found in common goldeneyes (Dow and Fredga 1985; Pöysä 1999; Pöysä 2006), common eiders (Fast et al. 2010; Lusignan et al. 2010), and red-breasted mergansers (Thimot et al. 2020). Interestingly, the only effect of our experimental treatments, when habitat features were controlled, was on the number of eggs laid and a trend towards fewer eggs hatching (Table 4, Figure 2). Perhaps nests with eggs attracted parasitic females to lay a few eggs in those nests, and nests closer and more directly facing the water might be more accessible. In any case, we did find that wood ducks readily used nest sites already containing eggs, suggesting that they do not avoid sites even where there is evidence of current occupancy. Odell and Eadie (2010) found a similar pattern in a separate experiment with wood ducks, suggesting that abandoned eggs could subsequently be included in a clutch of a new female who then incubates the nest. Whether this represents accidental “parasitism’ or more covertly, a form of “pre-emptive parasitism” is an intriguing question and may be a factor contributing to the high frequency of conspecific brood parasitism observed in this species.

Our results have management and conservation implications and offer some insight on the divergent trajectories that appear to characterize nest site selection studies by gamebird vs. non-gamebird bird ecologists. A large number of studies have now recognized the importance of both social and personal information use in nest site selection by birds (Ahlering et al. 2010; Ahlering and Faaborg 2006; Betts et al. 2008; Campobello and Sealy 2011; Chalfoun and Schmidt 2012; Coulton et al. 2011; Nocera and Betts 2010; Szymkowiak 2013; Ward et al. 2010). However, studies of waterfowl have focused more on habitat characteristics affecting nesting behaviors, while studies of passerines tend to focus more on the importance of social information (see also Eichholz and Elmberg 2014; O’Neil et al. 2014). Until now, the data on waterfowl have not been deeply integrated into the broader literature on public information use in other birds, but we advocate that both research realms would benefit by more cross-pollination (see O’Neil et al. 2014 for a similar perspective). The way different species and populations balance the use of personal versus social information undoubtedly varies, not only among species (the focus of much current literature), but also over different temporal and spatial scales. We suggest that the temporal scale of information use, in particular, has not been widely investigated – the types of cues used for initial nest site discovery might be very different from those used to refine or adjust nest selection decisions. Perhaps even more importantly, in light of both rapidly changing climates and habitats, and huge investments in habitat conservation, the pace at which each type of information varies could be critical (e.g., habitat structure is likely to change gradually whereas social cues related to reproductive success or performance could change drastically within a single nesting season; Betts et al. 2008). Wildlife biologists working to create or restore high quality nest site habitat may experience limited success if social cues are more important in the early stages of nest site discovery and attraction (Ahlering et al. 2010; Ahlering and Faaborg 2006; Nocera and Betts 2010; Ward et al. 2010). Conversely, providing evidence of social cues to attract birds to new habitats when the underlying habitat conditions are inadequate or deteriorating could attract birds into an ecological trap (Schlaepfer et al. 2002) unless birds use external habitat cues to avoid those locations initially. A deeper understanding of how multiple cues and sources of information are integrated throughout an individual’s lifetime and at critical life history junctures may have valuable conservation applications.

## Supporting information

Berg & Eadie 2020 Raw data file

## Declarations

### Ethics approval

All of our methods were approved by the UC Davis Institutional Animal Care and Use Committee (Protocol #15824). Females were caught, banded, measured, and released under a US Migratory Bird Banding (BBL) Master permit #10562 (to JME). All methods were observational.

### Authors’ contributions

ECB erected the nest boxes used in this study and collected the field data, with help from JME and student volunteers. JME conducted the statistical analyses. ECB and JME designed the study together and contributed equally to the writing of the manuscript.

## Acknowledgments

The authors would like to thank all of the students who helped with the field research, in particular Rick Ziegler and Richard Shaw. We also thank Tom Moore for help establishing the study site and for providing much feedback. Financial support for this project was provided by grants and fellowships from the University of California at Davis (to ECB), a National Science Foundation Research Training Grant in Animal Behavior (to ECB), a Sigma-Xi Grant-in-Aid of Research Award (to ECB), National Science Foundation Integrative Organismal Systems grant 1355208 (to JME), National Institute of Food and Agriculture grant WFB-6342-H (to JME), and funds from the Dennis G. Raveling Endowed Chair (to JME). The Putah Creek Council provided support for a field assistant, and the California Waterfowl Association donated funds for the wood duck boxes erected for this study. The manuscript was written with the help of sabbatical support from the American University of Paris (to ECB).

