## Supplementary material for "An experimental test of information use by North American wood ducks (*Aix sponsa*): external habitat cues, not social visual cues, influence initial nest-site selection": Berg & Eadie 2020 Raw data file

| Year | Site | Box | Erect | Charge date | Charge julian | Treat# | Treat | Use | #Nests | WODU use | Used Previoulsy? | Attempt | EndStage | 1stEggFound | First Egg Julian | HatchDate | Hatch Julian | #Eggs | #Hatch | BandDate | Recap | Minimum#Hens | Parasitic Eggs | Box Type | Height (m) |
|---|---|---|---|---|---|---|---|---|---|---|---|---|---|---|---|---|---|---|---|---|---|---|---|---|---|
| 1998 | PC | 1 | 15May1998 | 15May1998 | 135 | 1 | eggs | NA | 0 | Not Used | No | | | | | | | | | | | | | 1 | 3 |
| 1998 | PC | 2 | 15May1998 | 15May1998 | 135 | 1 | down | NA | 0 | Not Used | No | | | | | | | | | | | | | 1 | 3 |
| 1998 | PC | 3 | 15May1998 | 15May1998 | 135 | 1 | control | NA | 0 | Not Used | No | | | | | | | | | | | | | 1 | 3,2 |
| 1998 | PC | 4 | 13May1998 | 13May1998 | 133 | 1 | eggs | nest | 1 | Nest | No | a | hatch | 18May1998 | 138 | 22Jun1998 | 173 | 6 | 5 | 18Jun1998 | no | 1 | 0 | 1 | 3,4 |
| 1998 | PC | 5 | 17Apr1998 | 04May1998 | 124 | 1 | control | NA | 0 | Not Used | No | | | | | | | | | | | | | 1 | 3,5 |
| 1998 | PC | 6 | 17Apr1998 | 04May1998 | 124 | 1 | down | nest | 1 | Nest | No | a | hatch | 01Jun1998 | 152 | 28Jun1998 | 179 | 8 | 8 | 18Jun1998 | no | | 0 | 1 | 2,8 |
| 1998 | PC | 7 | 17Apr1998 | 04May1998 | 124 | | no | nest | 1 | Nest | No | a | hatch | 11May1998 | 131 | 12Jun1998 | 163 | 14 | 12 | | no | | 1 | 2 | 3,5 |
| 1998 | PC | 8 | 17Apr1998 | 04May1998 | 124 | 1 | down | nest | 1 | Nest | No | a | hatch | 11May1998 | 131 | 04Jul1998 | 185 | 12 | 12 | 03Jul1998 | no | 1 | 0 | 2 | 3,6 |
| 1998 | PC | 9 | 17Apr1998 | 04May1998 | 124 | 1 | control | PH | 0 | Other | No | | | | | | | | | | | | | 2 | 3,3 |
| 1998 | PC | 10 | 17Apr1998 | 04May1998 | 124 | 1 | eggs | NA | 0 | Not Used | No | | | | | | | | | | | | | 2 | 2,8 |
| 1998 | PC | 11 | 20Apr1998 | 04May1998 | 124 | 1 | eggs | NA | 0 | Not Used | No | | | | | | | | | | | | | 2 | 2,7 |
| 1998 | PC | 12 | 20Apr1998 | 04May1998 | 124 | 1 | control | PH | 0 | Other | No | | | | | | | | | | | | | 2 | 3,1 |
| 1998 | PC | 13 | 20Apr1998 | 04May1998 | 124 | 1 | down | NA | 0 | Not Used | No | | | | | | | | | | | | | 2 | 3 |
| 1998 | PC | 14 | 20Apr1998 | 04May1998 | 124 | 1 | eggs | NA | 0 | Not Used | No | | | | | | | | | | | | | 2 | 3 |
| 1998 | PC | 15 | 22Apr1998 | 04May1998 | 124 | 1 | down | NA | 0 | Not Used | No | | | | | | | | | | | | | 2 | 2,7 |
| 1998 | PC | 16 | 22Apr1998 | 04May1998 | 124 | 1 | control | NA | 0 | Not Used | No | | | | | | | | | | | | | 2 | 3,3 |
| 1998 | PC | 17 | 22Apr1998 | 06May1998 | 126 | 1 | eggs | NA | 0 | Not Used | No | | | | | | | | | | | | | 2 | 3,2 |
| 1998 | PC | 18 | 22Apr1998 | 06May1998 | 126 | 1 | control | nest | 1 | Nest | No | a | hatch | 18May1998 | 138 | 15Jun1998 | 166 | 9 | 9 | 03Jun1998 | no | | 0 | 2 | 3,5 |
| 1998 | PC | 19 | 24Apr1998 | 06May1998 | 126 | 1 | down | nest | 1 | Nest | No | a | hatch | 25May1998 | 145 | 30Jun1998 | 181 | 10 | 10 | 03Jun1998 | no | 1 | 0 | 2 | 3,3 |
| 1998 | PC | 20 | 24Apr1998 | 06May1998 | 126 | 1 | control | NA | 0 | Not Used | No | | | | | | | | | | | | | 2 | 2,9 |
| 1998 | PC | 21 | 24Apr1998 | 06May1998 | 126 | 1 | down | NA | 0 | Not Used | No | | | | | | | | | | | | | 2 | 2,5 |
| 1998 | PC | 22 | 29Apr1998 | 06May1998 | 126 | 1 | eggs | NA | 0 | Not Used | No | | | | | | | | | | | | | 2 | 2,8 |
| 1998 | PC | 23 | 29Apr1998 | 06May1998 | 126 | 1 | down | NA | 0 | Not Used | No | | | | | | | | | | | | | 2 | 2,7 |
| 1998 | PC | 24 | 24Apr1998 | 06May1998 | 126 | 1 | eggs | NA | 0 | Not Used | No | | | | | | | | | | | | | 2 | 3 |
| 1998 | PC | 25 | 24Apr1998 | 06May1998 | 126 | 1 | control | NA | 0 | Not Used | No | | | | | | | | | | | | | 2 | 2,7 |
| 1998 | PC | 26 | 27Apr1998 | 06May1998 | 126 | 1 | down | NA | 0 | Not Used | No | | | | | | | | | | | | | 2 | 3 |
| 1998 | PC | 27 | 27Apr1998 | 06May1998 | 126 | 1 | control | NA | 0 | Not Used | No | | | | | | | | | | | | | 2 | 3,2 |
| 1998 | PC | 28 | 27Apr1998 | 06May1998 | 126 | 1 | eggs | EUST | 0 | Other | No | | | | | | | | | | | | | 2 | 2,6 |
| 1998 | PC | 29 | 27Apr1998 | 08May1998 | 128 | 1 | eggs | rat | 0 | Other | No | | | | | | | | | | | | | 2 | 2,4 |
| 1998 | PC | 30 | | | | | no | nest | 1 | Nest | No | a | hatch | 06May1998 | 126 | 06Jun1998 | 157 | 12 | 12 | 21May1998 | no | | 0 | 2 | 2,8 |
| 1998 | PC | 31 | 29Apr1998 | 08May1998 | 128 | 1 | control | NA | 0 | Not Used | No | | | | | | | | | | | | | 2 | 3,5 |
| 1998 | PC | 32 | | 08May1998 | 128 | 1 | down | NA | 0 | Not Used | No | | | | | | | | | | | | | 2 | 2,4 |
| 1998 | PC | 33 | | | | | no | nest | 1 | Nest | No | a | hatch | 27Mar1998 | 86 | 24Apr1998 | 114 | 13 | 13 | 25Apr1998 | no | | 0 | 2 | 2,3 |
| 1998 | PC | 34 | 29Apr1998 | 08May1998 | 128 | 1 | control | NA | 0 | Not Used | No | | | | | | | | | | | | | 2 | 2,7 |
| 1998 | PC | 35 | | 01Mar1998 | 60 | 1 | eggs | NA | 0 | Not Used | No | | | | | | | | | | | | | 2 | 2,9 |
| 1998 | PC | 36 | 08May1998 | 08May1998 | 128 | 1 | down | NA | 0 | Not Used | No | | | | | | | | | | | | | 2 | 2,1 |
| 1998 | PC | 37 | 13May1998 | 13May1998 | 133 | 1 | control | NA | 0 | Not Used | No | | | | | | | | | | | | | 1 | 3 |
| 1999 | PC | 1 | 15May1998 | 07May1999 | 127 | | no | nest | 1 | Nest | No | a | hatch | 07May1999 | 127 | 11Jun1999 | 162 | 10 | 10 | 09Jun1999 | no | 2 | 0 | 1 | 3 |
| 1999 | PC | 2 | 15May1998 | 07May1999 | 127 | 1 | control | nest | 1 | Nest | No | a | hatch | 21May1999 | 141 | 24Jun1999 | 175 | 9 | 9 | 21Jun1999 | no | 1 | 0 | 1 | 3 |
| 1999 | PC | 3 | 15May1998 | 14Apr1999 | 104 | | no | nest | 1 | Nest | No | a | hatch | 14Apr1999 | 104 | 29Apr1999 | 119 | 16 | 10 | 14Apr1999 | no | | 3 | 1 | 3,2 |
| 1999 | PC | 3 | 15May1998 | 21May1999 | 141 | 1 | control | NA | 1 | Not Used | Yes | | | | | | | | | | | | | 1 | 3,2 |
| 1999 | PC | 4 | 13May1998 | 03Mar1999 | 62 | 1 | down | nest | 2 | Nest | Yes | a | laying | 02Apr1999 | 92 | | | 1 | 0 | | | 1 | 0 | 1 | 3,4 |
| 1999 | PC | 4 | 13May1998 | 03Mar1999 | 62 | 1 | down | nest | 2 | Nest | Yes | b | hatch | 23Apr1999 | 113 | 03Jun1999 | 154 | 15 | 14 | 24May1999 | no | 2 | 2 | 1 | 3,4 |
| 1999 | PC | 5 | 17Apr1998 | 03Mar1999 | 62 | 1 | eggs | nest | 1 | Nest | No | a | incubation | 17Mar1999 | 76 | | | 10 | 0 | | | 1 | 0 | 1 | 3,5 |
| 1999 | PC | 6 | 17Apr1998 | 03Mar1999 | 62 | 1 | control | nest | 1 | Nest | Yes | a | hatch | 07Apr1999 | 97 | 15May1999 | 135 | 12 | 10 | 12May1999 | no | 1 | 0 | 1 | 2,8 |
| 1999 | PC | 6 | 17Apr1998 | 21May1999 | 141 | 2 | down | EUST | 1 | Other | No | | | | | | | | | | | | | 1 | 2,8 |
| 1999 | PC | 7 | 17Apr1998 | 03Mar1999 | 62 | 1 | eggs | nest | 1 | Nest | Yes | a | laying | 26Mar1999 | 85 | 26Mar1999 | 85 | 4 | 0 | | | | 0 | 2 | 3,5 |
| 1999 | PC | 8 | 17Apr1998 | 03Mar1999 | 62 | 1 | down | nest | 1 | Nest | Yes | a | hatch | 26Mar1999 | 85 | 12May1999 | 132 | 17 | 15 | 23Apr1999 | no | 2 | 4 | 2 | 3,6 |
| 1999 | PC | 8 | 17Apr1998 | 21May1999 | 141 | 1 | eggs | EUST | 1 | Other | No | | | | | | | | | | | | | 2 | 3,6 |
| 1999 | PC | 9 | 17Apr1998 | 03Mar1999 | 62 | 1 | control | nest | 1 | Nest | No | a | hatch | 02Apr1999 | 92 | 26May1999 | 146 | 12 | 11 | 17May1999 | no | 1 | 0 | 2 | 3,3 |
| 1999 | PC | 10 | 17Apr1998 | 03Mar1999 | 62 | 1 | control | NA | 1 | Not Used | No | | | | | | | | | | | | | 2 | 2,8 |
| 1999 | PC | 10 | 17Apr1998 | 07May1999 | 127 | 2 | eggs | nest | 1 | Nest | Yes | a | hatch | 14May1999 | 134 | 26Jun1999 | 177 | 9 | 7 | 11Jun1999 | no | 1 | 0 | 2 | 2,8 |
| 1999 | PC | 11 | 20Apr1998 | 07Mar1999 | 66 | 1 | down | EUST | 0 | Other | No | | | | | | | | | | | | | 2 | 2,7 |
| 1999 | PC | 11 | 20Apr1998 | 07May1999 | 127 | 2 | down | NA | 0 | Not Used | No | | | | | | | | | | | | | 2 | 2,7 |
| 1999 | PC | 12 | 20Apr1998 | 07Mar1999 | 66 | 1 | eggs | EUST | 0 | Other | No | | | | | | | | | | | | | 2 | 3,1 |
| 1999 | PC | 12 | 20Apr1998 | 07May1999 | 127 | 2 | down | NA | 0 | Not Used | No | | | | | | | | | | | | | 2 | 3,1 |
| 1999 | PC | 13 | 20Apr1998 | 07Mar1999 | 66 | 1 | control | EUST | 0 | Other | No | | | | | | | | | | | | | 2 | 3 |
| 1999 | PC | 13 | 20Apr1998 | 07May1999 | 127 | 2 | control | NA | 0 | Not Used | No | | | | | | | | | | | | | 2 | 3 |
| 1999 | PC | 14 | 20Apr1998 | 07Mar1999 | 66 | 1 | eggs | NA | 0 | Not Used | No | | | | | | | | | | | | | 2 | 3 |
| 1999 | PC | 14 | 20Apr1998 | 07May1999 | 127 | 2 | eggs | NA | 0 | Not Used | No | | | | | | | | | | | | | 2 | 3 |
| 1999 | PC | 15 | 22Apr1998 | 07Mar1999 | 66 | 1 | down | NA | 0 | Not Used | No | | | | | | | | | | | | | 2 | 2,7 |
| 1999 | PC | 15 | 22Apr1998 | 07May1999 | 127 | 2 | control | NA | 0 | Not Used | No | | | | | | | | | | | | | 2 | 2,7 |
| 1999 | PC | 16 | 22Apr1998 | 07Mar1999 | 66 | 1 | down | SQ | 0 | Other | No | | | | | | | | | | | | | 2 | 3,3 |

Berg Eadie 2020: "An experimental test of information use by North American wood ducks (Aix sponsa): external habitat cues, not social visual cues, influence initial nest-site selection"
Raw data file

| | | | | | | | | | | | | | | | | | | | | | | | | | |
|---|---|---|---|---|---|---|---|---|---|---|---|---|---|---|---|---|---|---|---|---|---|---|---|---|---|
| 1999 | PC | 16 | 22Apr1998 | 07May1999 | 127 | 2 | eggs | NA | 0 | Not Used | No | | | | | | | | | | | | | 2 | 3,3 |
| 1999 | PC | 17 | 22Apr1998 | 07Mar1999 | 66 | 1 | control | nest | 1 | Nest | No | a | hatch | 02Apr1999 | 92 | 10May1999 | 130 | 7 | 7 | 03Jun1998 | yes | | 0 | 2 | 3,2 |
| 1999 | PC | 17 | 22Apr1998 | 21May1999 | 141 | 2 | control | NA | 1 | Not Used | Yes | | | | | | | | | | | | | 2 | 3,2 |
| 1999 | PC | 18 | 22Apr1998 | 07Mar1999 | 66 | 1 | eggs | SQ | 0 | Other | Yes | | | | | | | | | | | | | 2 | 3,5 |
| 1999 | PC | 18 | 22Apr1998 | 07May1999 | 127 | 2 | down | NA | 0 | Not Used | No | | | | | | | | | | | | | 2 | 3,5 |
| 1999 | PC | 19 | 24Apr1998 | 17Mar1999 | 76 | 1 | down | NA | 0 | Not Used | Yes | | | | | | | | | | | | | 2 | 3,3 |
| 1999 | PC | 19 | 24Apr1998 | 07May1999 | 127 | 2 | control | NA | 0 | Not Used | No | | | | | | | | | | | | | 2 | 3,3 |
| 1999 | PC | 20 | 24Apr1998 | 07Mar1999 | 66 | | no | WSOW | 0 | Other | No | | | | | | | | | | | | | 2 | 2,9 |
| 1999 | PC | 20 | 24Apr1998 | 07May1999 | 127 | 1 | down | NA | 0 | Not Used | No | | | | | | | | | | | | | 2 | 2,9 |
| 1999 | PC | 21 | 24Apr1998 | 07Mar1999 | 66 | 1 | eggs | WSOW | 0 | Other | No | | | | | | | | | | | | | 2 | 2,5 |
| 1999 | PC | 22 | 29Apr1998 | 07Mar1999 | 66 | 1 | control | nest | 1 | Nest | No | a | hatch | 12Apr1999 | 102 | 23May1999 | 143 | 10 | 1 | 19May1999 | no | 1 | 0 | 2 | 2,8 |
| 1999 | PC | 23 | 29Apr1998 | 07Mar1999 | 66 | 1 | down | SQ | 0 | Other | No | | | | | | | | | | | | | 2 | 2,7 |
| 1999 | PC | 23 | 29Apr1998 | 05May1999 | 125 | 2 | eggs | NA | 0 | Not Used | No | | | | | | | | | | | | | 2 | 2,7 |
| 1999 | PC | 24 | 24Apr1998 | 07Mar1999 | 66 | 1 | control | SQ | 0 | Other | No | | | | | | | | | | | | | 2 | 3 |
| 1999 | PC | 24 | 24Apr1998 | 05May1999 | 125 | 2 | down | NA | 0 | Not Used | No | | | | | | | | | | | | | 2 | 3 |
| 1999 | PC | 25 | 24Apr1998 | 07Mar1999 | 66 | 1 | eggs | NA | 0 | Not Used | No | | | | | | | | | | | | | 2 | 2,7 |
| 1999 | PC | 25 | 24Apr1998 | 05May1999 | 125 | 2 | control | NA | 0 | Not Used | No | | | | | | | | | | | | | 2 | 2,7 |
| 1999 | PC | 26 | 27Apr1998 | 07Mar1999 | 66 | 1 | control | nest | 1 | Nest | No | a | hatch | 21Mar1999 | 80 | 29Apr1999 | 119 | 13 | 11 | | no | 2 | 0 | 2 | 3 |
| 1999 | PC | 26 | 27Apr1998 | 21May1999 | 141 | 2 | down | NA | 1 | Not Used | Yes | | | | | | | | | | | | | 2 | 3 |
| 1999 | PC | 27 | 27Apr1998 | 07Mar1999 | 66 | | no | WSOW | 0 | Other | No | | | | | | | | | | | | | 2 | 3,2 |
| 1999 | PC | 28 | 27Apr1998 | 07Mar1999 | 66 | | no | WSOW | 0 | Other | No | | | | | | | | | | | | | 2 | 2,6 |
| 1999 | PC | 29 | 27Apr1998 | 07Mar1999 | 66 | 1 | down | EUST | 0 | Other | No | | | | | | | | | | | | | 2 | 2,4 |
| 1999 | PC | 29 | 27Apr1998 | 05May1999 | 125 | 2 | eggs | EUST | 0 | Other | No | | | | | | | | | | | | | 2 | 2,4 |
| 1999 | PC | 30 | | ? | | | no | nest | 1 | Nest | No | a | hatch | 07Mar1999 | 66 | 18Apr1999 | 108 | 5 | 5 | 14Apr1999 | yes | 1 | 0 | 2 | 2,8 |
| 1999 | PC | 30 | | 21May1999 | 141 | 1 | eggs | NA | 1 | Not Used | No | | | | | | | | | | | | | 2 | 2,8 |
| 1999 | PC | 31 | 29Apr1998 | 10Mar1999 | 69 | 1 | eggs | NA | 0 | Not Used | No | | | | | | | | | | | | | 2 | 3,5 |
| 1999 | PC | 31 | 29Apr1998 | 05May1999 | 125 | 2 | control | NA | 0 | Not Used | No | | | | | | | | | | | | | 2 | 3,5 |
| 1999 | PC | 32 | | 10Mar1999 | 69 | 1 | down | nest | 1 | Nest | No | a | hatch | 02Apr1999 | 92 | 12May1999 | 132 | 11 | 9 | 28Apr1999 | no | 1 | 0 | 2 | 2,4 |
| 1999 | PC | 32 | | 21May1999 | 141 | 2 | control | NA | 1 | Not Used | Yes | | | | | | | | | | | | | 2 | 2,4 |
| 1999 | PC | 33 | | ? | | 1 | eggs | nest | 1 | Nest | Yes | a | hatch | 28Mar1999 | 87 | 12May1999 | 132 | 13 | 12 | 25Apr1998 | yes | 3 | 0 | 2 | 2,3 |
| 1999 | PC | 33 | | 21May1999 | 141 | 2 | down | NA | 1 | Not Used | No | | | | | | | | | | | | | 2 | 2,3 |
| 1999 | PC | 34 | 29Apr1998 | 10Mar1999 | 69 | 1 | control | NA | 0 | Not Used | No | | | | | | | | | | | | | 2 | 2,7 |
| 1999 | PC | 34 | 29Apr1998 | 05May1999 | 125 | 2 | down | NA | 0 | Not Used | No | | | | | | | | | | | | | 2 | 2,7 |
| 1999 | PC | 35 | | ? | | 1 | control | nest | 1 | Nest | No | a | hatch | 10Mar1999 | 69 | 24Apr1999 | 114 | 13 | 11 | 25Apr1999 | no | 1 | 0 | 2 | 2,9 |
| 1999 | PC | 35 | | 21May1999 | 141 | 2 | eggs | NA | 1 | Not Used | Yes | | | | | | | | | | | | | 2 | 2,9 |
| 1999 | PC | 36 | 08May1998 | 10Mar1999 | 69 | 1 | down | EUST | 0 | Other | No | | | | | | | | | | | | | 2 | 2,1 |
| 1999 | PC | 36 | 08May1998 | 07May1999 | 127 | 2 | eggs | NA | 0 | Not Used | No | | | | | | | | | | | | | 2 | 2,1 |
| 1999 | PC | 37 | 13May1998 | 10Mar1999 | 69 | 1 | eggs | NA | 0 | Not Used | No | | | | | | | | | | | | | 1 | 3 |
| 1999 | PC | 37 | 13May1998 | 07May1999 | 127 | 2 | down | NA | 0 | Not Used | No | | | | | | | | | | | | | 1 | 3 |
| 1999 | RR | 1 | | 13Mar1999 | 72 | 1 | control | nest | 1 | Nest | No | a | hatch | 26Apr1999 | 116 | 01Jun1999 | 152 | 10 | 10 | 01Jun1999 | no | | 0 | 1 | 3 |
| 1999 | RR | 2 | | 13Mar1999 | 72 | 1 | eggs | nest | 1 | Nest | No | a | hatch | 26Apr1999 | 116 | 01Jun1999 | 152 | 15 | 9 | 01Jun1999 | no | 1 | 2 | 1 | 2,8 |
| 1999 | RR | 3 | | 13Mar1999 | 72 | 1 | down | NA | 1 | Not Used | No | | | | | | | | | | | | | 2 | 2,5 |
| 1999 | RR | 3 | | 05May1999 | 125 | 2 | control | nest | 1 | Nest | No | a | hatch | 19May1999 | 139 | 19Jun1999 | 170 | 11 | 11 | 09Jun1999 | no | 1 | 0 | 2 | 2,5 |
| 1999 | RR | 4 | | 13Mar1999 | 72 | 1 | control | nest | 1 | Nest | No | a | hatch | 03Apr1999 | 93 | 13May1999 | 133 | 11 | 11 | 28Apr1999 | no | 1 | 0 | 1 | 2,5 |
| 1999 | RR | 4 | | 24May1999 | 144 | 2 | down | NA | 1 | Not Used | Yes | | | | | | | | | | | | | 1 | 2,5 |
| 1999 | RR | 5 | | 13Mar1999 | 72 | 1 | down | nest | 1 | Nest | No | a | hatch | 28Mar1999 | 87 | 08May1999 | 128 | 11 | 11 | | yes | 1 | 0 | 2 | 3 |
| 1999 | RR | 6 | | | | | no | WSOW | 0 | Other | No | | | | | | | | | | | | | 1 | 2,8 |
| 1999 | RR | 6 | | 24May1999 | 144 | 2 | control | NA | 0 | Not Used | No | | | | | | | | | | | | | 1 | 2,8 |
| 1999 | RR | 7 | | 13Mar1999 | 72 | 1 | eggs | WSOW | 0 | Other | No | | | | | | | | | | | | | 2 | 2,8 |
| 1999 | RR | 7 | | 24May1999 | 144 | 2 | eggs | NA | 0 | Not Used | No | | | | | | | | | | | | | 2 | 2,8 |
| 1999 | RR | 8 | | 14Mar1999 | 73 | 1 | down | NA | 0 | Not Used | No | | | | | | | | | | | | | 2 | 2,9 |
| 1999 | RR | 8 | | 05May1999 | 125 | 2 | eggs | NA | 0 | Not Used | No | | | | | | | | | | | | | 2 | 2,9 |
| 1999 | RR | 9 | | | | 1 | control | nest | 2 | Nest | No | a | hatch | 14Mar1999 | 73 | 21Apr1999 | 111 | 14 | 10 | 19Apr1999 | no | 1 | 1 | 2 | 2,5 |
| 1999 | RR | 9 | | 24May1999 | 144 | 2 | control | nest | 2 | Nest | Yes | b | | 02Jun1999 | 153 | 6/9 - 7/16 | | 9 | 9 | 19Apr1999 | yes | | 0 | 2 | 2,5 |
| 1999 | RR | 10 | | 14Mar1999 | 73 | 1 | control | nest | 1 | Nest | No | a | hatch | 02Apr1999 | 92 | 11May1999 | 131 | 9 | 9 | 21Apr1999 | no | 1 | 0 | 1 | 2,5 |
| 1999 | RR | 10 | | 24May1999 | 144 | 2 | down | NA | 1 | Not Used | Yes | | | | | | | | | | | | | 1 | 2,5 |
| 1999 | RR | 11 | | 14Mar1999 | 73 | 1 | eggs | nest | 1 | Nest | No | a | hatch | 02Apr1999 | 92 | 11May1999 | 131 | 7 | 7 | 28Apr1999 | no | 1 | 0 | 2 | 2,6 |
| 1999 | RR | 11 | | 24May1999 | 144 | 2 | eggs | NA | 1 | Not Used | Yes | | | | | | | | | | | | | 2 | 2,6 |
| 1999 | RR | 12 | | 14Mar1999 | 73 | 1 | control | PH | 1 | Other | No | | | | | | | | | | | | | 2 | 2,5 |
| 1999 | RR | 12 | | 05May1999 | 125 | 2 | down | nest | 1 | Nest | No | a | hatch | 12May1999 | 132 | 21Jun1999 | 172 | 12 | 9 | 09Jun1999 | no | 1 | 0 | 2 | 2,5 |

Berg Eadie 2020: "An experimental test of information use by North American wood ducks (Aix sponsa): external habitat cues, not social visual cues, influence initial nest-site selection"
Raw data file

| Dist. Water (m) | Dir. Water(Degrees) | Tree Species | Comp Dir. (Degrees) | Tree 0 (m) | Tree 90 (m) | Tree 180 (m) | Tree 270 (m) | % Vis | %Cov G | % Cov E | %Cov 5m | Std Type | Std Size | Std Density | Canopy Ht (m) | Slp (degrees) | % Shcov | East (m) | North (m) | zone | Dist next box (m) |
|---|---|---|---|---|---|---|---|---|---|---|---|---|---|---|---|---|---|---|---|---|---|
| 8 | 180 | EUCA | 300 | 150 | 12 | 0 | 4 | 40 | 20 | 60 | 40 | EUCA | 3 | 1 | 12 | 15 | 5 | 609434,66 | 4263995,64 | 10 | 82,93 |
| 20 | 180 | EUCA | 334 | 2 | 3 | 4 | 2 | 0 | 25 | 50 | 60 | EUCA | 4 | 3 | 15 | 10 | 15 | 609353,12 | 4263983,6 | 10 | 130,2 |
| 2 | 270 | EUCA | 260 | 150 | 2 | 4 | 6 | 40 | 80 | 60 | 80 | EUCA | 4 | 2 | 12 | 45 | 35 | 609224,63 | 4263964,84 | 10 | 320,5 |
| 3 | 90 | EUCA | 44 | 6 | 10 | 2,5 | 12 | 10 | 50 | 50 | 60 | MIX | 4 | 2 | 12 | 30 | 5 | 609903,66 | 4263966,62 | 10 | 238,7 |
| 15 | 180 | EUCA | 340 | 0 | 3 | 5 | 4 | 45 | 20 | 80 | 90 | JUCA | 4 | 3 | 13 | 20 | 10 | 608665,4 | 4263962,9 | 10 | 231,8 |
| 15 | 270 | EUCA | 242 | 30 | 30 | 10 | 20 | 10 | 60 | 5 | 20 | EUCA | 0 | 0 | 15 | 10 | 0 | 608433,5 | 4263961 | 10 | 279,2 |
| 40 | 90 | EUCA | 33 | 11 | 4 | 3 | 8 | 0 | 50 | 95 | 25 | EUCA | 0 | 0 | 11 | 10 | 5 | 608154,43 | 4263973,28 | 10 | 262,9 |
| 13 | 180 | EUCA | 19 | 35 | 8 | 13 | 12 | 40 | 10 | 75 | 60 | EUCA | 0 | 0 | 8 | 25 | 0 | 607892,45 | 4263997,43 | 10 | 292,5 |
| 20 | 180 | EUCA | 343 | 50 | 55 | 15 | 13 | 5 | 20 | 70 | 35 | EUCA | 0 | 0 | 7 | 30 | 0 | 607600,38 | 4263977,93 | 10 | 222,2 |
| 55 | 270 | POFE | 297 | 70 | 50 | 45 | 25 | 0 | 80 | 90 | 90 | POFE | 0 | 0 | 8 | 20 | 0 | 607379,29 | 4263997,55 | 10 | 425 |
| 15 | 180 | JUCA | 29 | 150 | 15 | 7 | 4 | 0 | 100 | 80 | 90 | JUCA | 1 | 1 | 10 | 45 | 0 | 606977,91 | 4264137,97 | 10 | 57,61 |
| 20 | 270 | JUCA | 319 | 150 | 6 | 8 | 5 | 0 | 100 | 85 | 85 | JUCA | 3 | 2 | 13 | 45 | 0 | 606925,53 | 4264164,85 | 10 | 59,43 |
| 35 | 180 | QULO | 4 | 150 | 20 | 10 | 11 | 0 | 90 | 85 | 85 | QULO | 0 | 0 | 8 | 45 | 0 | 606873,5 | 4264195,8 | 10 | 163,3 |
| 45 | 180 | JUCA | 348 | 150 | 40 | 15 | 35 | 0 | 100 | 95 | 85 | JUCA | 0 | 0 | 7 | 30 | 20 | 606738,62 | 4264286,55 | 10 | 120,6 |
| 60 | 90 | JUCA | 60 | 8 | 12 |  | 13 | 0 | 95 | 100 | 90 | JUCA | 1 | 0 | 7 | 30 | 0 | 606636,56 | 4264351 | 10 | 57,5 |
| 20 | 270 | EUCA | 314 | 10 | 25 | 9 | 4 | 0 | 80 | 15 | 20 | MIX | 2 | 2 | 12 | 15 | 40 | 606581,32 | 4264364,08 | 10 | 126,8 |
| 30 | 180 | JUCA | 47 | 150 | 10 | 15 | 9 | 0 | 80 | 95 | 95 | JUCA | 1 | 1 | 8 | 35 | 0 | 606485,33 | 4264446,5 | 10 | 144,7 |
| 20 | 180 | EUCA | 29 | 10 | 13 | 1 | 2 | 0 | 85 | 80 | 80 | MIX | 4 | 3 | 10 | 10 | 30 | 606357,83 | 4264518,08 | 10 | 41,54 |
| 5 | 180 | QULO | 76 | 1 | 1 | 4 | 3 | 50 | 95 | 60 | 75 | MIX | 4 | 2 | 9 | 25 | 55 | 606321,44 | 4264538,22 | 10 | 41,07 |
| 10 | 180 | QULO | 342 | 1 | 2 | 1 |  | 0 | 100 | 15 | 60 | MIX | 4 | 2 | 12 | 20 | 45 | 606286,56 | 424559,81 | 10 | 45,04 |
| 18 | 270 | EUCA | 316 | 150 | 3 | 5 | 20 | 0 | 100 | 30 | 40 | MIX | 1 | 1 | 12 | 20 | 5 | 606252,43 | 4264590,97 | 10 | 56,68 |
| 13 | 180 | EUCA | 30 |  | 1 | 2 |  | 0 | 100 | 75 | 85 | EUCA | 3 | 3 | 13 | 20 | 30 | 606198,06 | 4264601 | 10 | 69,34 |
| 18 | 180 | SALX | 4 | 5 | 2 | 5 | 1,5 | 0 | 90 | 70 | 80 | MIX | 4 | 2 | 8 | 5 | 15 | 606129,52 | 4264617 | 10 | 86,54 |
| 8 | 270 | QULO | 218 | 5 | 4 |  | 2 | 0 | 90 | 60 | 60 | MIX | 4 | 3 | 12 | 15 | 75 | 606044,76 | 4264599,72 | 10 | 51,68 |
| 13 | 270 | SALX | 262 | 2 | 5 | 4 | 5 | 0 | 100 | 50 | 55 | SALX | 4 | 2 | 9 | 15 | 25 | 605992,62 | 4264602,68 | 10 | 89,84 |
| 6 | 270 | JUCA | 274 | 8 | 4 | 4 | 4 | 20 | 75 | 60 | 75 | MIX | 3 | 2 | 9 | 30 | 10 | 605891,23 | 4264580,57 | 10 | 138,2 |
| 12 | 180 | EUCA | 2 | 150 | 5 | 4 | 5 | 5 | 80 | 75 | 55 | MIX | 4 | 2 | 7 | 20 | 25 | 605754,07 | 4264601,63 | 10 | 469 |
| 11 | 180 | QULO | 349 | 150 | 10 | 11 | 10 | 30 | 100 | 75 | 80 | QULO | 1 | 0 | 11 | 45 | 0 | 605346,33 | 4264836,35 | 10 | 137,6 |
| 9 | 180 | QULO | 30 | 150 | 20 | 7 | 10 | 5 | 100 | 90 | 85 | QULO | 1 | 1 | 8 | 45 | 0 | 605228,85 | 4264908,61 | 10 | 419,9 |
| 18 | 180 | QULO | 353 | 4 | 18 | 6 | 6 | 30 | 85 | 50 | 40 | MIX | 2 | 1 | 15 | 45 | 5 | 604837,56 | 4265060,4 | 10 | 115,1 |
| 13 | 180 | EUCA | 354 | 2 | 1 | 4 | 2 | 0 | 85 | 80 | 80 | EUCA | 4 | 4 | 15 | 10 | 55 | 604723,35 | 4265048 | 10 | 87,44 |
| 8 | 180 | POFE | 349 | 6 | 4 | 7 | 3 | 30 | 100 | 40 | 60 | EUCA | 4 | 2 | 9 | 15 | 45 | 604639,3 | 4265019,8 | 10 | 81,27 |
| 11 | 270 | JUCA | 274 | 18 | 4 | 1,5 | 2 | 10 | 75 | 60 | 85 | MIX | 4 | 1 | 10 | 20 | 15 | 604558,36 | 4265019,74 | 10 | 42,45 |
| 8 | 270 | EUCA | 251 | 10 | 5 |  | 6 | 10 | 75 | 50 | 50 | MIX | 2 | 1 | 13 | 25 | 0 | 604517,31 | 4265009,51 | 10 | 135,68 |
| 10 | 270 | EUCA | 288 | 12 | 6 | 4 | 5 | 0 | 90 | 30 | 40 | EUCA | 3 | 2 | 14 | 20 | 25 | 604382,68 | 4264992,88 | 10 | 152,5 |
| 9 | 270 | EUCA | 310 | 150 | 9 | 6 | 6 | 0 | 85 | 50 | 45 | MIX | 4 | 2 | 13 | 15 | 45 | 604229,94 | 4264976,11 | 10 | 104,8 |
| 5 | 180 | EUCA | 28 | 4 | 7 | 3 | 6 | 30 | 100 | 75 | 80 | EUCA | 4 | 2 | 16 | 30 | 15 | 604124,84 | 4264967,66 | 10 |  |
| 8 | 180 | EUCA | 300 | 150 | 12 | 0 | 4 | 40 | 20 | 60 | 40 | EUCA | 3 | 1 | 12 | 15 | 5 | 609434,66 | 4263995,64 | 10 | 82,93 |
| 20 | 180 | EUCA | 334 | 2 | 3 | 4 | 2 | 0 | 25 | 50 | 60 | EUCA | 4 | 3 | 15 | 10 | 15 | 609353,12 | 4263983,6 | 10 | 130,2 |
| 2 | 270 | EUCA | 260 | 150 | 2 | 4 | 6 | 40 | 80 | 60 | 80 | EUCA | 4 | 2 | 12 | 45 | 35 | 609224,63 | 4263964,84 | 10 | 320,5 |
| 2 | 270 | EUCA | 260 | 150 | 2 | 4 | 6 | 40 | 80 | 60 | 80 | EUCA | 4 | 2 | 12 | 45 | 35 | 609224,63 | 4263964,84 | 10 | 320,5 |
| 3 | 90 | EUCA | 44 | 6 | 10 | 2,5 | 12 | 10 | 50 | 50 | 60 | MIX | 4 | 2 | 12 | 30 | 5 | 609903,66 | 4263966,62 | 10 | 238,7 |
| 3 | 90 | EUCA | 44 | 6 | 10 | 2,5 | 12 | 10 | 50 | 50 | 60 | MIX | 4 | 2 | 12 | 30 | 5 | 609903,66 | 4263966,62 | 10 | 238,7 |
| 15 | 180 | EUCA | 340 | 0 | 3 | 5 | 4 | 45 | 20 | 80 | 90 | JUCA | 4 | 3 | 13 | 20 | 10 | 608665,4 | 4263962,9 | 10 | 231,8 |
| 15 | 270 | EUCA | 242 | 30 | 30 | 10 | 20 | 10 | 60 | 5 | 20 | EUCA | 0 | 0 | 15 | 10 | 0 | 608433,5 | 4263961 | 10 | 279,2 |
| 15 | 270 | EUCA | 242 | 30 | 30 | 10 | 20 | 10 | 60 | 5 | 20 | EUCA | 0 | 0 | 15 | 10 | 0 | 608433,5 | 4263961 | 10 | 279,2 |
| 40 | 90 | EUCA | 33 | 11 | 4 | 3 | 8 | 0 | 50 | 95 | 25 | EUCA | 0 | 0 | 11 | 10 | 5 | 608154,43 | 4263973,28 | 10 | 262,9 |
| 13 | 180 | EUCA | 19 | 35 | 8 | 13 | 12 | 40 | 10 | 75 | 60 | EUCA | 0 | 0 | 8 | 25 | 0 | 607892,45 | 4263997,43 | 10 | 292,5 |
| 13 | 180 | EUCA | 19 | 35 | 8 | 13 | 12 | 40 | 10 | 75 | 60 | EUCA | 0 | 0 | 8 | 25 | 0 | 607892,45 | 4263997,43 | 10 | 292,5 |
| 20 | 180 | EUCA | 343 | 50 | 55 | 15 | 13 | 5 | 20 | 70 | 35 | EUCA | 0 | 0 | 7 | 30 | 0 | 607600,38 | 4263977,93 | 10 | 222,2 |
| 55 | 270 | POFE | 297 | 70 | 50 | 45 | 25 | 0 | 80 | 90 | 90 | POFE | 0 | 0 | 8 | 20 | 0 | 607379,29 | 4263997,55 | 10 | 425 |
| 55 | 270 | POFE | 297 | 70 | 50 | 45 | 25 | 0 | 80 | 90 | 90 | POFE | 0 | 0 | 8 | 20 | 0 | 607379,29 | 4263997,55 | 10 | 425 |
| 15 | 180 | JUCA | 29 | 150 | 15 | 7 | 4 | 0 | 100 | 80 | 90 | JUCA | 1 | 1 | 10 | 45 | 0 | 606977,91 | 4264137,97 | 10 | 57,61 |
| 15 | 180 | JUCA | 29 | 150 | 15 | 7 | 4 | 0 | 100 | 80 | 90 | JUCA | 1 | 1 | 10 | 45 | 0 | 606977,91 | 4264137,97 | 10 | 57,61 |
| 20 | 270 | JUCA | 319 | 150 | 6 | 8 | 5 | 0 | 100 | 85 | 85 | JUCA | 3 | 2 | 13 | 45 | 0 | 606925,53 | 4264164,85 | 10 | 59,43 |
| 20 | 270 | JUCA | 319 | 150 | 6 | 8 | 5 | 0 | 100 | 85 | 85 | JUCA | 3 | 2 | 13 | 45 | 0 | 606925,53 | 4264164,85 | 10 | 59,43 |
| 35 | 180 | QULO | 4 | 150 | 20 | 10 | 11 | 0 | 90 | 85 | 85 | QULO | 0 | 0 | 8 | 45 | 0 | 606873,5 | 4264195,8 | 10 | 163,3 |
| 35 | 180 | QULO | 4 | 150 | 20 | 10 | 11 | 0 | 90 | 85 | 85 | QULO | 0 | 0 | 8 | 45 | 0 | 606873,5 | 4264195,8 | 10 | 163,3 |
| 45 | 180 | JUCA | 348 | 150 | 40 | 15 | 35 | 0 | 100 | 95 | 85 | JUCA | 0 | 0 | 7 | 30 | 20 | 606738,62 | 4264286,55 | 10 | 120,6 |
| 45 | 180 | JUCA | 348 | 150 | 40 | 15 | 35 | 0 | 100 | 95 | 85 | JUCA | 0 | 0 | 7 | 30 | 20 | 606738,62 | 4264286,55 | 10 | 120,6 |
| 60 | 90 | JUCA | 60 | 8 | 12 |  | 13 | 0 | 95 | 100 | 90 | JUCA | 1 | 0 | 7 | 30 | 0 | 606636,56 | 4264351 | 10 | 57,5 |
| 60 | 90 | JUCA | 60 | 8 | 12 |  | 13 | 0 | 95 | 100 | 90 | JUCA | 1 | 0 | 7 | 30 | 0 | 606636,56 | 4264351 | 10 | 57,5 |
| 20 | 270 | EUCA | 314 | 10 | 25 | 9 | 4 | 0 | 80 | 15 | 20 | MIX | 2 | 2 | 12 | 15 | 40 | 606581,32 | 4264364,08 | 10 | 126,8 |

Berg Eadie 2020: "An experimental test of information use by North American wood ducks (Aix sponsa): external habitat cues, not social visual cues, influence initial nest-site selection"
Raw data file

| | | | | | | | | | | | | | | | | | | | | | |
|---|---|---|---|---|---|---|---|---|---|---|---|---|---|---|---|---|---|---|---|---|---|
| 20 | 270 | EUCA | 314 | 10 | 25 | 9 | 4 | 0 | 80 | 15 | 20 | MIX | 2 | 2 | 12 | 15 | 40 | 606581,32 | 4264364,08 | 10 | 126,8 |
| 30 | 180 | JUCA | 47 | 150 | 10 | 15 | 9 | 0 | 80 | 95 | 95 | JUCA | 1 | 1 | 8 | 35 | 0 | 606485,33 | 4264446,5 | 10 | 144,7 |
| 30 | 180 | JUCA | 47 | 150 | 10 | 15 | 9 | 0 | 80 | 95 | 95 | JUCA | 1 | 1 | 8 | 35 | 0 | 606485,33 | 4264446,5 | 10 | 144,7 |
| 20 | 180 | EUCA | 29 | 10 | 13 | 1 | 2 | 0 | 85 | 80 | 80 | MIX | 4 | 3 | 10 | 10 | 30 | 606357,83 | 4264518,08 | 10 | 41,54 |
| 20 | 180 | EUCA | 29 | 10 | 13 | 1 | 2 | 0 | 85 | 80 | 80 | MIX | 4 | 3 | 10 | 10 | 30 | 606357,83 | 4264518,08 | 10 | 41,54 |
| 5 | 180 | QULO | 76 | 1 | 1 | 4 | 3 | 50 | 95 | 60 | 75 | MIX | 4 | 2 | 9 | 25 | 55 | 606321,44 | 4264538,22 | 10 | 41,07 |
| 5 | 180 | QULO | 76 | 1 | 1 | 4 | 3 | 50 | 95 | 60 | 75 | MIX | 4 | 2 | 9 | 25 | 55 | 606321,44 | 4264538,22 | 10 | 41,07 |
| 10 | 180 | QULO | 342 | 1 | 2 | 1 | | 0 | 100 | 15 | 60 | MIX | 4 | 2 | 12 | 20 | 45 | 606286,56 | 424559,81 | 10 | 45,04 |
| 10 | 180 | QULO | 342 | 1 | 2 | 1 | | 0 | 100 | 15 | 60 | MIX | 4 | 2 | 12 | 20 | 45 | 606286,56 | 424559,81 | 10 | 45,04 |
| 18 | 270 | EUCA | 316 | 150 | 3 | 5 | 20 | 0 | 100 | 30 | 40 | MIX | 1 | 1 | 12 | 20 | 5 | 606252,43 | 4264590,97 | 10 | 56,68 |
| 13 | 180 | EUCA | 30 | | 1 | 2 | | 0 | 100 | 75 | 85 | EUCA | 3 | 3 | 13 | 20 | 30 | 606198,06 | 4264601 | 10 | 69,34 |
| 18 | 180 | SALX | 4 | 5 | 2 | 5 | 1,5 | 0 | 90 | 70 | 80 | MIX | 4 | 2 | 8 | 5 | 15 | 606129,52 | 4264617 | 10 | 86,54 |
| 18 | 180 | SALX | 4 | 5 | 2 | 5 | 1,5 | 0 | 90 | 70 | 80 | MIX | 4 | 2 | 8 | 5 | 15 | 606129,52 | 4264617 | 10 | 86,54 |
| 8 | 270 | QULO | 218 | 5 | 4 | | 2 | 0 | 90 | 60 | 60 | MIX | 4 | 3 | 12 | 15 | 75 | 606044,76 | 4264599,72 | 10 | 51,68 |
| 8 | 270 | QULO | 218 | 5 | 4 | | 2 | 0 | 90 | 60 | 60 | MIX | 4 | 3 | 12 | 15 | 75 | 606044,76 | 4264599,72 | 10 | 51,68 |
| 13 | 270 | SALX | 262 | 2 | 5 | 4 | 5 | 0 | 100 | 50 | 55 | SALX | 4 | 2 | 9 | 15 | 25 | 605992,62 | 4264602,68 | 10 | 89,84 |
| 13 | 270 | SALX | 262 | 2 | 5 | 4 | 5 | 0 | 100 | 50 | 55 | SALX | 4 | 2 | 9 | 15 | 25 | 605992,62 | 4264602,68 | 10 | 89,84 |
| 6 | 270 | JUCA | 274 | 8 | 4 | 4 | 4 | 20 | 75 | 60 | 75 | MIX | 3 | 2 | 9 | 30 | 10 | 605891,23 | 4264580,57 | 10 | 138,2 |
| 6 | 270 | JUCA | 274 | 8 | 4 | 4 | 4 | 20 | 75 | 60 | 75 | MIX | 3 | 2 | 9 | 30 | 10 | 605891,23 | 4264580,57 | 10 | 138,2 |
| 12 | 180 | EUCA | 2 | 150 | 5 | 4 | 5 | 5 | 80 | 75 | 55 | MIX | 4 | 2 | 7 | 20 | 25 | 605754,07 | 4264601,63 | 10 | 469 |
| 11 | 180 | QULO | 349 | 150 | 10 | 11 | 10 | 30 | 100 | 75 | 80 | QULO | 1 | 0 | 11 | 45 | 0 | 605346,33 | 4264836,35 | 10 | 137,6 |
| 9 | 180 | QULO | 30 | 150 | 20 | 7 | 10 | 5 | 100 | 90 | 85 | QULO | 1 | 1 | 8 | 45 | 0 | 605228,85 | 4264908,61 | 10 | 419,9 |
| 9 | 180 | QULO | 30 | 150 | 20 | 7 | 10 | 5 | 100 | 90 | 85 | QULO | 1 | 1 | 8 | 45 | 0 | 605228,85 | 4264908,61 | 10 | 419,9 |
| 18 | 180 | QULO | 353 | 4 | 18 | 6 | 6 | 30 | 85 | 50 | 40 | MIX | 2 | 1 | 15 | 45 | 5 | 604837,56 | 4265060,4 | 10 | 115,1 |
| 18 | 180 | QULO | 353 | 4 | 18 | 6 | 6 | 30 | 85 | 50 | 40 | MIX | 2 | 1 | 15 | 45 | 5 | 604837,56 | 4265060,4 | 10 | 115,1 |
| 13 | 180 | EUCA | 354 | 2 | 1 | 4 | 2 | 0 | 85 | 80 | 80 | EUCA | 4 | 4 | 15 | 10 | 55 | 604723,35 | 4265048 | 10 | 87,44 |
| 13 | 180 | EUCA | 354 | 2 | 1 | 4 | 2 | 0 | 85 | 80 | 80 | EUCA | 4 | 4 | 15 | 10 | 55 | 604723,35 | 4265048 | 10 | 87,44 |
| 8 | 180 | POFE | 349 | 6 | 4 | 7 | 3 | 30 | 100 | 40 | 60 | EUCA | 4 | 2 | 9 | 15 | 45 | 604639,3 | 4265019,8 | 10 | 81,27 |
| 8 | 180 | POFE | 349 | 6 | 4 | 7 | 3 | 30 | 100 | 40 | 60 | EUCA | 4 | 2 | 9 | 15 | 45 | 604639,3 | 4265019,8 | 10 | 81,27 |
| 11 | 270 | JUCA | 274 | 18 | 4 | 1,5 | 2 | 10 | 75 | 60 | 85 | MIX | 4 | 1 | 10 | 20 | 15 | 604558,36 | 4265019,74 | 10 | 42,45 |
| 11 | 270 | JUCA | 274 | 18 | 4 | 1,5 | 2 | 10 | 75 | 60 | 85 | MIX | 4 | 1 | 10 | 20 | 15 | 604558,36 | 4265019,74 | 10 | 42,45 |
| 8 | 270 | EUCA | 251 | 10 | 5 | | 6 | 10 | 75 | 50 | 50 | MIX | 2 | 1 | 13 | 25 | 0 | 604517,31 | 4265009,51 | 10 | 135,68 |
| 8 | 270 | EUCA | 251 | 10 | 5 | | 6 | 10 | 75 | 50 | 50 | MIX | 2 | 1 | 13 | 25 | 0 | 604517,31 | 4265009,51 | 10 | 135,68 |
| 10 | 270 | EUCA | 288 | 12 | 6 | 4 | 5 | 0 | 90 | 30 | 40 | EUCA | 3 | 2 | 14 | 20 | 25 | 604382,68 | 4264992,88 | 10 | 152,5 |
| 10 | 270 | EUCA | 288 | 12 | 6 | 4 | 5 | 0 | 90 | 30 | 40 | EUCA | 3 | 2 | 14 | 20 | 25 | 604382,68 | 4264992,88 | 10 | 152,5 |
| 9 | 270 | EUCA | 310 | 150 | 9 | 6 | 6 | 0 | 85 | 50 | 45 | MIX | 4 | 2 | 13 | 15 | 45 | 604229,94 | 4264976,11 | 10 | 104,8 |
| 9 | 270 | EUCA | 310 | 150 | 9 | 6 | 6 | 0 | 85 | 50 | 45 | MIX | 4 | 2 | 13 | 15 | 45 | 604229,94 | 4264976,11 | 10 | 104,8 |
| 5 | 180 | EUCA | 28 | 4 | 7 | 3 | 6 | 30 | 100 | 75 | 80 | EUCA | 4 | 2 | 16 | 30 | 15 | 604124,84 | 4264967,66 | 10 | |
| 5 | 180 | EUCA | 28 | 4 | 7 | 3 | 6 | 30 | 100 | 75 | 80 | EUCA | 4 | 2 | 16 | 30 | 15 | 604124,84 | 4264967,66 | 10 | |
| 15 | 270 | QULO | 268 | 8 | 2 | 1 | 2 | 40 | 95 | 70 | 70 | EUCA | 2 | 1 | 10 | | 70 | 597392,25 | 4266206,98 | 10 | 115,2 |
| 12 | 270 | QULO | 302 | 4 | 8 | 5 | 4 | 15 | 95 | 20 | 40 | QULO | 4 | 2 | 12 | 50 | 55 | 597494,87 | 4266155,13 | 10 | 81,45 |
| 13 | 180 | POFE | 15 | 2 | 1,5 | 4 | 4 | 0 | 100 | 60 | 75 | MIX | 3 | 3 | 12 | 30 | 40 | 597575,42 | 4266124,65 | 10 | 116,7 |
| 13 | 180 | POFE | 15 | 2 | 1,5 | 4 | 4 | 0 | 100 | 60 | 75 | MIX | 3 | 3 | 12 | 30 | 40 | 597575,42 | 4266124,65 | 10 | 116,7 |
| 11 | 180 | JUCA | 12 | 7 | 3 | 5 | 8 | 0 | 98 | 80 | 95 | MIX | 2 | 2 | 10 | 40 | 30 | 597693,61 | 4266124,24 | 10 | 117,8 |
| 11 | 180 | JUCA | 12 | 7 | 3 | 5 | 8 | 0 | 98 | 80 | 95 | MIX | 2 | 2 | 10 | 40 | 30 | 597693,61 | 4266124,24 | 10 | 117,8 |
| 10 | 180 | QULO | 300 | 5 | 6 | 5 | 9 | 40 | 95 | 65 | 65 | MIX | 4 | 2 | 10 | 45 | 20 | 597808,82 | 4266120,41 | 10 | 111,7 |
| 7 | 270 | JUCA | 290 | 6 | 6 | 0 | 5 | 35 | 100 | 80 | 80 | MIX | 3 | 2 | 10 | 60 | 75 | 597923,1 | 4266121,24 | 10 | 103 |
| 7 | 270 | JUCA | 290 | 6 | 6 | 0 | 5 | 35 | 100 | 80 | 80 | MIX | 3 | 2 | 10 | 60 | 75 | 597923,1 | 4266121,24 | 10 | 103 |
| 20 | 270 | QULO | 254 | 5 | 15 | 10 | 1 | 5 | 100 | 75 | 75 | MIX | 3 | 2 | 11 | 30 | 5 | 598022,52 | 4266148,66 | 10 | 202,3 |
| 20 | 270 | QULO | 254 | 5 | 15 | 10 | 1 | 5 | 100 | 75 | 75 | MIX | 3 | 2 | 11 | 30 | 5 | 598022,52 | 4266148,66 | 10 | 202,3 |
| 18 | 180 | JUCA | 324 | 10 | 2,5 | 8 | 3 | 0 | 100 | 90 | 90 | JUCA | 4 | 2 | 11 | 20 | 0 | 598218,32 | 4266198,14 | 10 | 115,1 |
| 18 | 180 | JUCA | 324 | 10 | 2,5 | 8 | 3 | 0 | 100 | 90 | 90 | JUCA | 4 | 2 | 11 | 20 | 0 | 598218,32 | 4266198,14 | 10 | 115,1 |
| 5 | 180 | JUCA | 18 | 11 | 5 | 1 | 9 | 20 | 95 | 75 | 70 | MIX | 2 | 1 | 11 | 50 | 0 | 598332,86 | 4266188,06 | 10 | 140,7 |
| 5 | 180 | JUCA | 18 | 11 | 5 | 1 | 9 | 20 | 95 | 75 | 70 | MIX | 2 | 1 | 11 | 50 | 0 | 598332,86 | 4266188,06 | 10 | 140,7 |
| 17 | 180 | QULO | 9 | 2 | 3 | 10 | 2 | 40 | 95 | 80 | 80 | MIX | 1 | 4 | 8 | 30 | 0 | 598471,26 | 4266168,07 | 10 | 116,5 |
| 17 | 180 | QULO | 9 | 2 | 3 | 10 | 2 | 40 | 95 | 80 | 80 | MIX | 1 | 4 | 8 | 30 | 0 | 598471,26 | 4266168,07 | 10 | 116,5 |
| 7 | 90 | QULO | 60 | 3 | 5 | 4 | 3 | 5 | 100 | 80 | 85 | MIX | 3 | 2 | 13 | 30 | 25 | 598575,67 | 4266126,55 | 10 | 207,4 |
| 7 | 90 | QULO | 60 | 3 | 5 | 4 | 3 | 5 | 100 | 80 | 85 | MIX | 3 | 2 | 13 | 30 | 25 | 598575,67 | 4266126,55 | 10 | 207,4 |
| 10 | 180 | POFE | 318 | 5 | 2 | 7 | 1 | 30 | 90 | 55 | 45 | POFE | 3 | 2 | 11 | 35 | 70 | 598778,51 | 426067,73 | 10 | |
| 10 | 180 | POFE | 318 | 5 | 2 | 7 | 1 | 30 | 90 | 55 | 45 | POFE | 3 | 2 | 11 | 35 | 70 | 598778,51 | 426067,73 | 10 | |
